# Differential polarization of cortical pyramidal neuron dendrites through weak extracellular fields

**DOI:** 10.1101/216184

**Authors:** Florian Aspart, Michiel W.H. Remme, Klaus Obermayer

## Abstract

The rise of transcranial current stimulation (tCS) techniques have sparked an increasing interest in the effects of weak extracellular electric fields on neural activity. These fields modulate ongoing neural activity through polarization of the neuronal membrane. While the somatic polarization has been investigated experimentally, the frequency-dependent polarization of the dendritic trees in the presence of alternating (AC) fields has received little attention yet. Using a biophysically detailed model with experimentally constrained active conductances, we analyze the subthreshold response of cortical pyramidal cells to weak AC fields, as induced during tCS. We observe a strong frequency resonance around 10-20 Hz in the apical dendrites sensitivity to polarize in response to electric fields but not in the basal dendrites nor the soma. To disentangle the relative roles of the cell morphology and active and passive membrane properties in this resonance, we perform a thorough analysis using simplified models, e.g. a passive pyramidal neuron model, simple passive cables and reconstructed cell model with simplified ion channels. We attribute the origin of the resonance in the apical dendrites to (i) a locally increased sensitivity due to the morphology and to (ii) the high density of h-type channels. Our systematic study provides an improved understanding of the subthreshold response of cortical cells to weak electric fields and, importantly, allows for an improved design of tCS stimuli.

## Introduction

In the past decade, evidence of ephaptic coupling [1,2](see [3] for a review) and the dawn of transcranial current stimulation (tCS) have sparked an increasing interest in the modulation of neuronal activity through weak extracellular fields. tCS can modulate ongoing neural activity [4] and cognitive capacity, e.g. to potentiate memory [5], and its non-invasiveness provides it with a great potential for clinical applications. However, the design of optimized stimulation strategies is impeded by the poor understanding of tCS’s mechanism of action on neural tissue.

tCS makes use of the application of low amplitude direct (tDCS) or alternating (tACS) current on the scalp. This current induces a weak extracellular electric field within the brain [6]. Its typically low amplitude (≤ 2V/m) is comparable to that of endogenous fields [1]. Unlike the latter, fields due to tCS can be considered as spatially uniform at the cell scale [7]. Weak uniform electric field have been shown to induce inward and outward membrane currents at different cell locations [8]. This results in a non-uniform membrane polarization of the neuron (in the order of 0.1 mV per V/m) [9]. While the resulting somatic polarization alters spike timing of the neuron [10, 11], terminal polarization, e.g. dendritic, modulates synaptic efficacy [12,13]. Experimental and theoretical work demonstrated that the somatic polarization is stronger for fields orientated parallel to the somato-dendritic axis, depends on cell morphology [8] and decreases with the field frequency [14]. However, little is known about the frequency-dependence of the polarization in the dendritic tree, which is where virtually all synaptic input to a neuron arrives and which is endowed with many active, voltage-dependent processes [15] that might be modulated by extracellular fields.

In the present modelling study, we investigate the role of cell morphology and active membrane properties on the frequency-dependent polarization due to weak electric fields. More specifically, we consider the sensitivity of a biophysical cortical pyramidal cell model to become polarized in response to a weak electric field parallel to its somato-dendritic axis. We first consider the model with reconstructed morphology but without active channels and explain the main characteristics of its response using a simple passive cable model. We then consider the pyramidal cell model with active conductances. We find a strong frequency resonance in its field sensitivity at the apical dendrite which we relate to both the cells passive properties and the presence of specific ion channels. Finally, we use a simplified model of active conductances to generalize the relation between the cells active properties and its sensitivity to sinusoidal electric fields.

## Results

### Polarization of a passive pyramidal neuron in response to electric fields

#### Field sensitivity of a passive pyramidal neuron

To begin with, we consider the morphologically reconstructed layer 5b pyramidal neuron model published by *Hay et al.* [16] without active membrane properties. We simulate the subthreshold response of this model to an extracellular electric field, i.e. a gradient of extracellular potential. We choose the field orientation parallel to the cell’s apical dendrites. In the present work, we restrict ourselves to fields induced during transcranial stimulation. These fields are spatially uniform, i.e. their spatial derivative is null, and of low amplitude (1 V/m) [7]. We consider only Direct Current (DC) fields, *E*(*x,t*) = *E*_0_, or Alternating Current (AC), *E*(*x,t*) = *E*_0_ sin(2*πƒ*_*t*_*t*) where *ƒ*_*t*_ is the field frequency (see Methods). Such fields are known to polarize the membrane potential of the cell in a morphology-dependent manner [8,9]. When subject to a Direct Current (DC) field, the apical dendrites get oppositely polarized compared to the soma and basal dendrites (Fig. 1ABC). The polarization amplitude is strongest at the apical end. At the field onset, the basal and somatic regions display a slight overshoot. Due to the passive model’s linearity, the polarization is reversed when the field has opposite sign.

**Figure 1.**
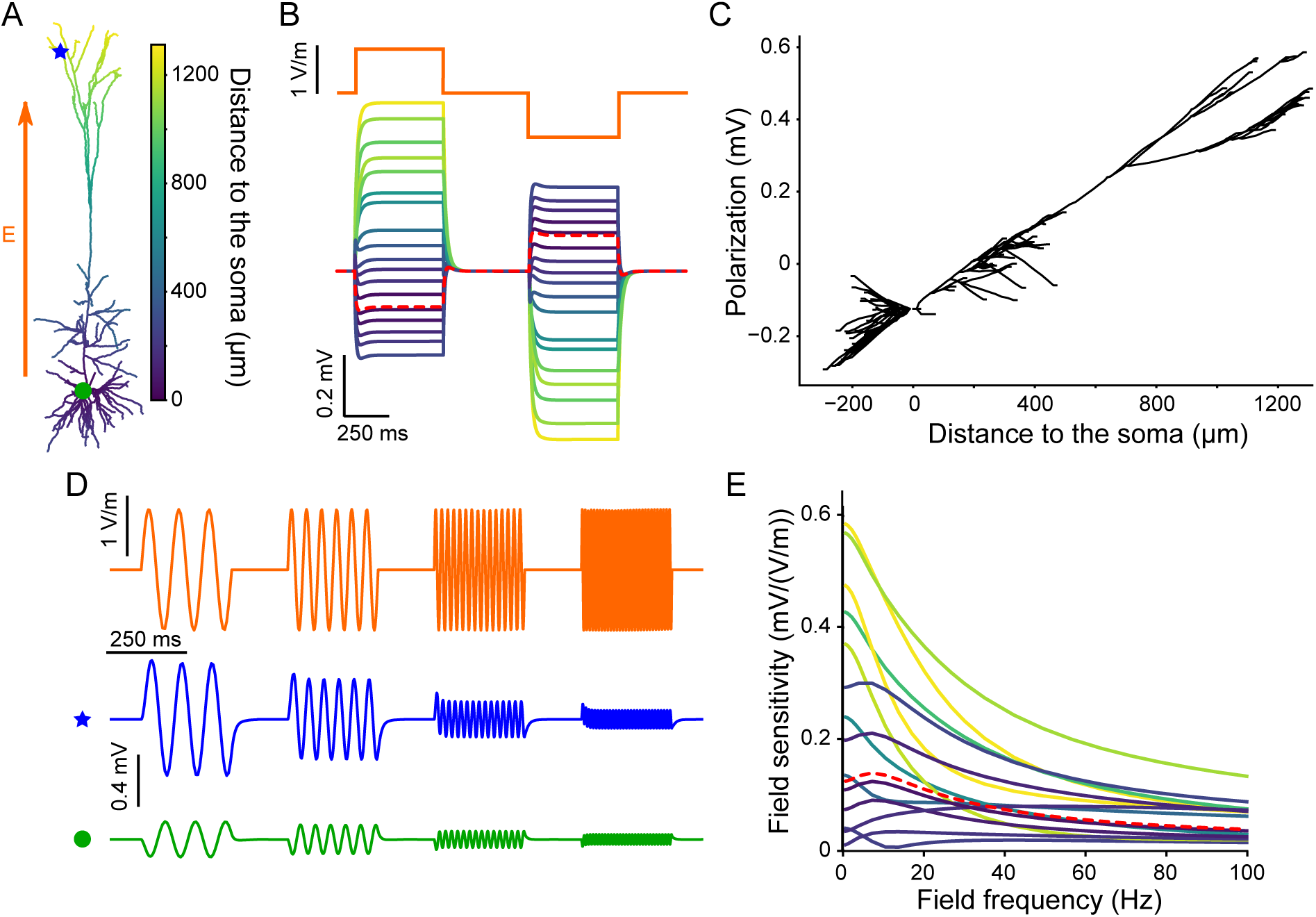
In a passive pyramidal cell model subject to an electric field, soma and basal dendrites get oppositely polarized than apical dendrites. This later are the most responsive to low frequency stimulation. (A) Considered neuron morphology with color coding the distance of each segment to the soma. (B) (Bottom) Membrane polarization of the passive cell due to positive and negative steps of DC electric field (orange, top). (C) Polarization due to a positive 1 V/m field plotted as the function of the distance from the soma. For clarity basal dendrites are plotted with negative distance. (D) Example polarization at the apical dendrite (blue star, bottom) and soma (green circle, middle) due to an an oscillating field of diverse frequencies (orange, top). (E) Frequency-dependent sensitivity of different cell segments to AC fields. Colors of the polarization (B) and field sensitivity (E) correspond to the distance from the soma as depicted in A. The red dashed lines correspond to the soma.

We further consider AC fields, which induce a sinusoidal polarization of the membrane (Fig.1D). The cell being passive, the polarization amplitude scales linearly with the field amplitude. Here and in the following, we define the field sensitivity of the cell at a given location as the ratio between the polarization amplitude and the field amplitude [14]. For most of the considered locations, the field sensitivity decreases with frequency (Fig. 1E). Compared to the basal dendrites and the soma, the apical dendrite is more sensitive to low frequency fields but its field sensitivity decreases faster with frequency. In the basal dendrites and the soma, the field sensitivity displays a slight frequency resonance around 10 Hz. Interestingly, some basal and proximal apical (oblique) branches show a strong frequency resonance from 20Hz to 50Hz (see Fig. S1 in Supporting information). This atypical field sensitivity profile corresponds to branches whose tip, when projected on the field axis, is further away from the soma than their branching point (see below). Nevertheless, the peak field sensitivity at this resonance is rather low compared to the maximal sensitivity observed for very slow fields at other locations.

In summary, the apical dendrites of our passive pyramidal neuron model are more sensitive to low frequency AC fields than the soma and basal dendrites. The field sensitivity decreases with frequency but at some proximal dendrites a resonance appears. Until the end of this section, we use a simpler model, namely a passive cable model, to investigate the origins of (i) the field sensitivity differences between the apical dendrites and the soma/basal dendrites and of (ii) the passive resonance.

#### Sensitivity of a passive cable to an extracellular field

We now consider a passive dendritic cable subject to an extracellular electric field *E* parallel to the cable axis, 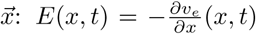, where *v*_*e*_ is the extracellular potential. Note that in this paragraph we do not limit ourselves to spatially uniform field but also consider non-uniform fields. The membrane potential *v*(*x,t*), i.e. the difference between intra-and extracellular potentials, is the solution to the cable equation [17]:

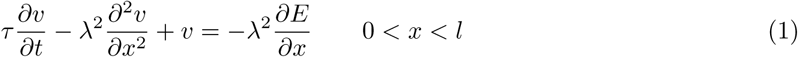

with the boundary conditions (sealed-end):

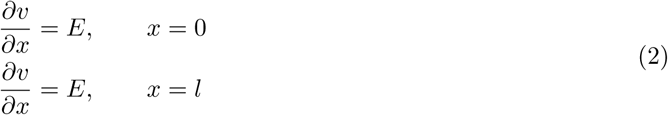

where *l*, *τ* and *λ* are respectively the cable’s length, the membrane time constant and space constant. Assuming the spatial and temporal components of the extracellular potential are independent, i.e. *v*_*e*_(*x,t*) = *v*_*x,e*_(*x*)*v*_*t,e*_(*t*), we solve Eq. 1 for arbitrary electric fields (see Methods for the details on the full derivation). Note that, under this assumption, the field is equivalent to correlated input currents distributed along the cable (right hand side, rhs, of Eq. 1) and at the extremities (rhs of Eq. 2).

AC fields induce a temporally sinusoidal polarization along the cable, whose amplitude scales linearly with the field amplitude since the cable is passive. We express the polarization as:

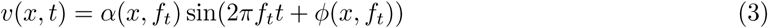

where *α*(*x*, *ƒ*_*t*_) and *φ*(*x*, *ƒ*_*t*_) are the polarization amplitude and its phase shift relative to the field. The sensitivity to the electric field of a neuron model, as defined for the reconstructed cell model, is equal to:

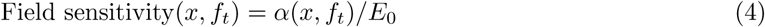

Hence, at a given frequency, the field sensitivity along the cable depends on the cable’s membrane time constant *τ*, its electrotonic length scale (or space constant) *λ*, and its length *l*. To further reduce the parameter space, we normalize the length units and define the cable electrotonic length, *L* = *l*/*λ* (in *λ*), and the electrotonic field amplitude, *E*_*λ*_ = *E*_0_/*λ* (in *V/λ*). Consequently, we express the field sensitivity in *V/(V/λ*).

The space constant *λ* represents the distance, in an infinite cable, over which membrane polarization due to a steady state input current decreases by 1/*e* [18]. In case of AC input, we can define a generalized frequency-dependent space constant *λ*_*gen*_(*ƒ*_*t*_) as:

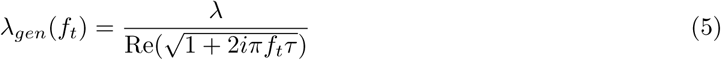

where Re(*z*) is the real part of the complex number *z*. As a consequence, high frequency inputs have more local effects than low frequency inputs [18].

Translated to the electric field, this implies that the response to higher frequency electric fields depends more strongly, in case of non-uniform field, on their local profile than for low frequency perturbations. In the following, we illustrate the practical implications of this effect for straight and bent cables and relate it to our observations in the reconstructed cell.

#### Field sensitivity of straight cables

In case of a straight cable subject to a spatially uniform field, the polarization at both ends of the cable is antisymmetric, i.e. they get oppositely polarized with the same amplitude (see Fig. 2A). Due to the field spatial uniformity, the effects of the field (rhs of Eq. 1) along the cable are null and the field is equivalent to correlated inputs at the extremities (Eq. 2). These inputs are of equivalent amplitude and opposite sign, more specifically inward at one end and outward at the other, which causes the antisymmetric polarization. For all frequencies, the sensitivity to AC fields is maximum at the end points, decreases toward the middle of the cable where a phase shift occurs, and then increases again (Fig. 2B). For low frequency, the field sensitivity is null solely at the cable center due to its antisymmetric nature. For very high frequency, e.g. *ƒ*_*t*_ > 200Hz, a broader area around the center shows virtually no polarization by the field. This area is less affected by the cable extremities due to the low frequency-dependent space constant, *λ*_*gen*_(*ƒ*_*t*_). This is locally equivalent to an infinite cable, which is not affected by uniform field.

**Figure 2.**
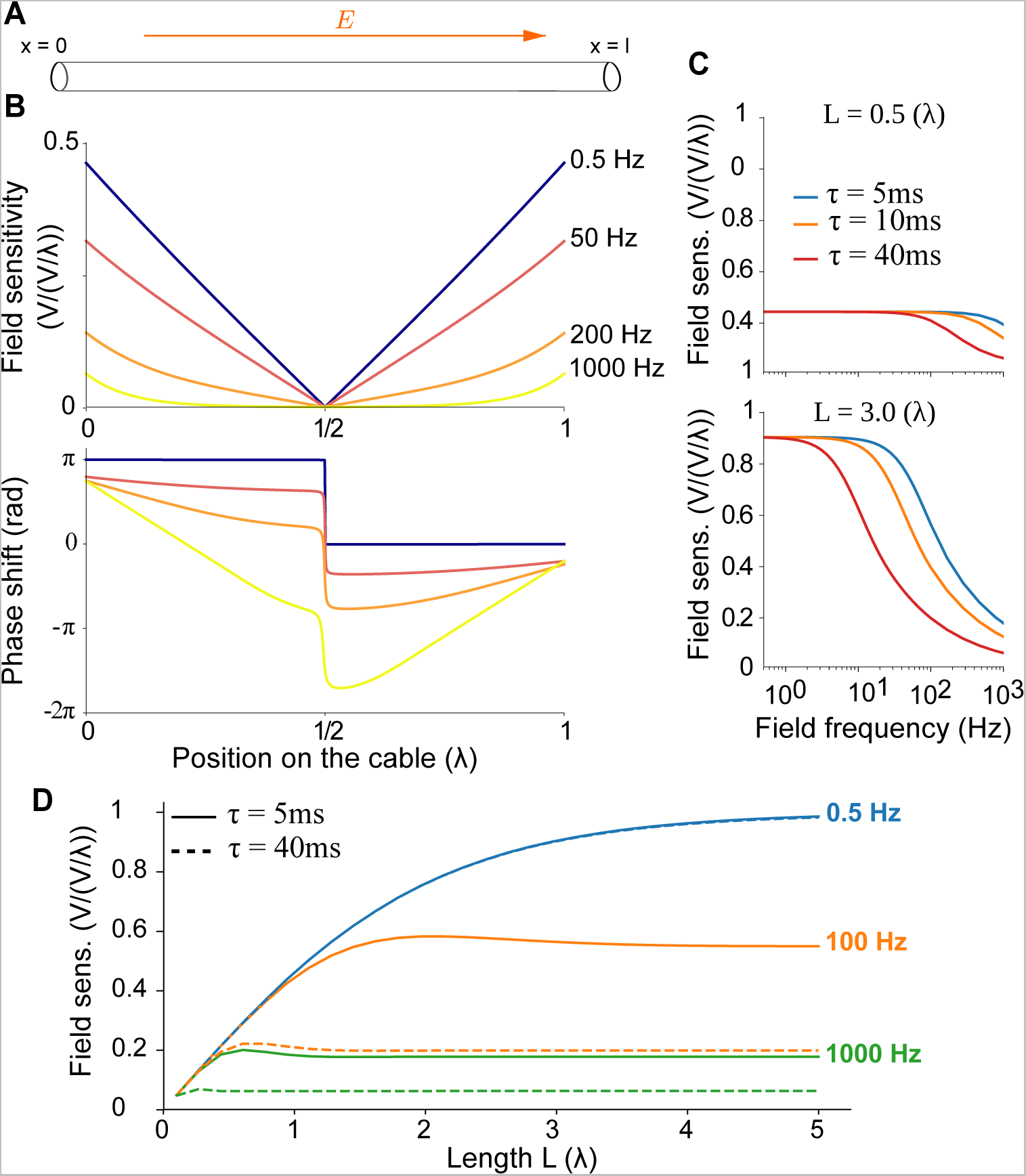
The polarization of a passive straight cable due to an extracellular field is anti-symmetric and decreases with the field frequency. (A) Schematic representation of a straight cable subject to a parallel electric field. The variables are detailed in Methods. (B) (top) Field sensitivity, i.e. ratio between membrane polarization and field amplitude, along the cable for different field frequencies (0.5, 50, 200 and 1000Hz, color coded) for τ = 40*ms* and *L* = 1λ. (bottom) Phase shift between the field and the membrane polarization oscillations. (C) Field sensitivity at the cable ends as a function of frequency (x axes), for various cable’s electrotonic length *L* (top: 0.5λ, bottom: 3λ) and time constant *τ* (color coded). (D) Field sensitivity at the cable ends as a function of electrotonic length L, for various frequencies (color coded) and membrane time constants (solid lines: 5ms, dashed lines: 40ms).

Overall, the longer the cable, the higher is the sensitivity to low frequency fields at the cable extremities (Fig. 2D). Intuitively, the field induces currents flowing inward at one cable end and outward at the other (Eq. 2). The polarization at both cable ends is opposite and cancels out in the middle. With increasing cable length, the distance separating these end points increases and their interaction decreases. As a result, the field sensitivity increases until it saturates when one end no longer affects the other end, and the cable can locally be considered as semi-infinite.

For all considered parameters, the field sensitivity decreases with frequency. The cutoff frequency is determined by the membrane time constant *τ* and the electrotonic length L (Fig. 2C). The higher the time constant, the slower the membrane reacts to perturbations and the faster the field sensitivity decays with frequency. At a given frequency and time constant *τ*, the sensitivity is independent of the total electrotonic cable length, L, provided the cable is long enough to be considered as semi-infinite at its end (Fig. 2D).

#### Field sensitivity of bent cables

In the presence of a uniform field, the straight cable model displays an opposite polarization at both ends and its sensitivity decreases with field frequency. However, it fails to reproduce some properties of the reconstructed cell model field sensitivity, such as the stronger polarization at the apical dendrites or the resonance effect in some proximal dendrites. To investigate the role of the morphology in these phenomena, we now consider a bent cable, as depicted in Fig. 3A, subject to a uniform extracellular field. We refer to the branch parallel to the field as the main part, the bent part being the branch with an angle of Θ compared to the field axis. This setup is similar to having a non-uniform extracellular field, where the extracellular potential spatial component is computed by projecting the bent part of the cable on the field axis:

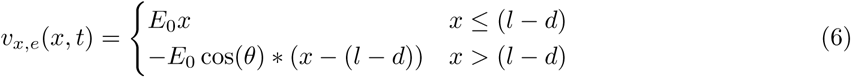

where *d* is the length of the bent branch, *x* ∈ [0, *l*] the position on the cable, 0 and *l* being respectively the extremities of the main and bent branches. Using this projection and analytical expression for the polarization of a straight cable through a field of arbitrary spatial profile, we compute the sensitivity to a spatially uniform field at both ends of the bent cable.

**Figure 3.**
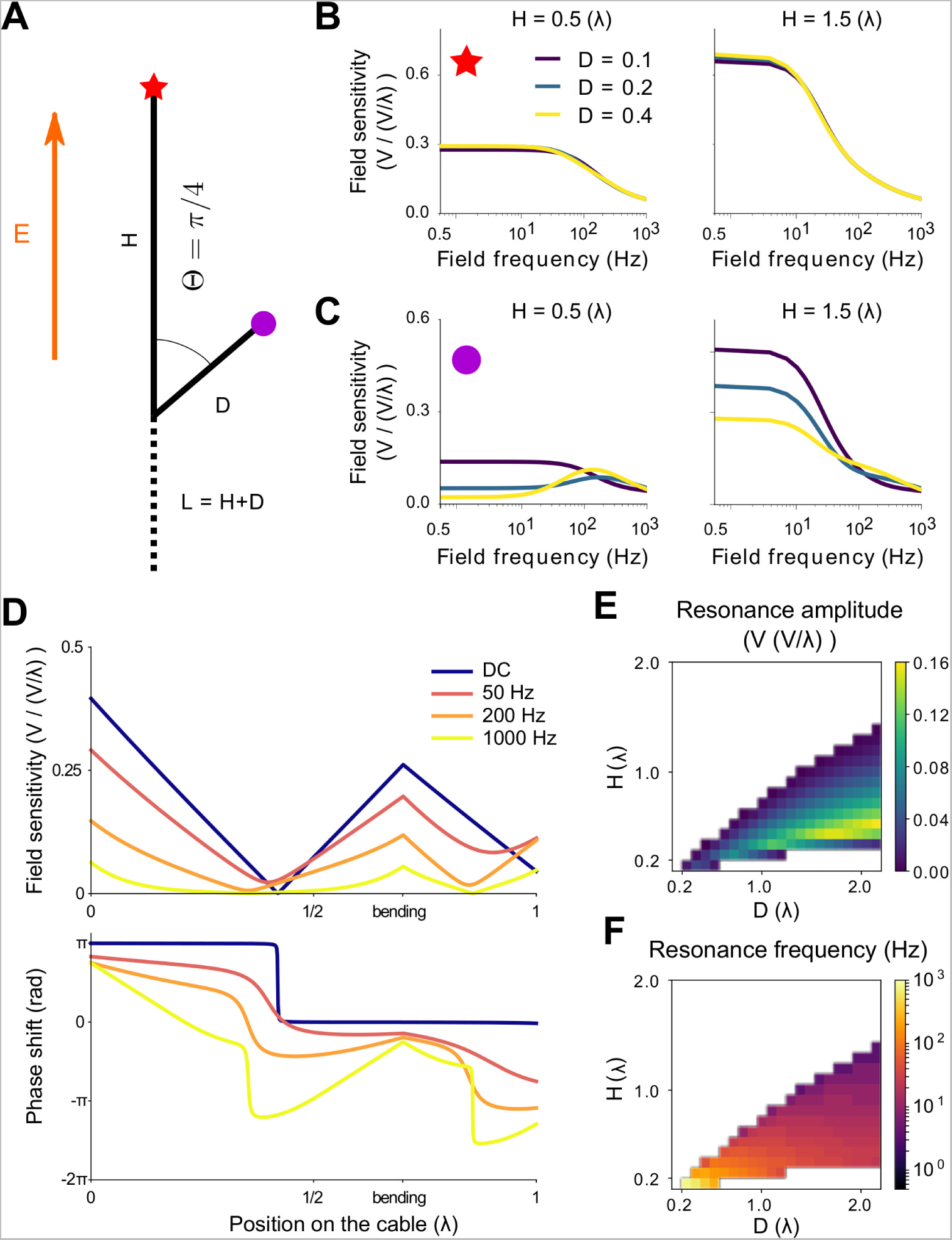
Passive cable with an acute bending angle can display a resonance in their sensitivity to spatially uniform fields. (A) Schematic representation of a bent cable. The unbent branch of the cable, of length *H*, is parallel to the field axis. The bent branch, of length *D*, have an angle Θ with the field. *L* is the total cable length. (B,C) Sensitivity ( in V/(V/λ)) at both cable ends, i.e. at the main (B) and bent (C) branches ends, as function of the field frequency. The sensitivities are displayed for various main (*H*, columns) and bent (*D* color coded) branches lengths. (D) Distribution of the sensitivity (top) and phase (bottom) along the bent cable for different field frequencies (0.5, 50, 200 and 1000Hz) for *H* = 0.6(λ) and *D* = 0.4(λ). (E) Resonance amplitude (sensitivity at the resonance minus sensitivity at 0.5Hz) and (F) resonance frequency at the bent end depending on both branches length. The white area corresponds to the absence of resonance. In all the plots the bending angle is Θ = *π*/4 (rad) and the membrane time constant *τ* = 40(*ms*). *L, H* and *D* are electrotonic lengths.

Compared to the straight cable case, the field sensitivity depends on two additional parameters: the normalized length (“electrotonic length”) of the bent branch, *D* = *d*/λ, and the bending angle Θ. The cable is no longer symmetric and the polarization differs at both ends. For obtuse angles (Θ ≥ *π*/2), the bending does not qualitatively modify the frequency-dependent sensitivity profile at both ends (Fig. S2 in Supporting Information). For example, if Θ = 3*π*/4, the field sensitivity at the bent end decreases with the length of the bent branch (for constant cable length), while the field sensitivity at the opposite extremity is little affected. This explains the different polarization amplitude in the apical and basal dendrite in the reconstructed pyramidal model. The basal dendrites and the soma correspond to the bent branch: there are more branches growing radially to the field. On the contrary, the main part of the cable represents the apical dendrites. These are more parallel to the field axis, e.g. the somato-dendritic axis, and have a higher sensitivity to low frequency fields.

Interestingly, for acute bending angles (Θ < *π*/2), the field sensitivity profile at the end point of the bent branch changes qualitatively (see Fig. 3B for Θ = *π*/4). At this location, the sensitivity to low frequency fields decreases with the length of the bent branch and a resonance appears. The range of frequencies affected by this decrease, and therefore the resonance frequency, is determined by the membrane time constant *τ*: the larger the membrane time constant, the lower the cut-off frequency (Fig. S4 in Supporting Information). At the end point of the main branch, the field sensitivity decreases with the length of the bending *D*, unless *L* is sufficiently long. For fixed *D*, it keeps a similar dependence on the remaining parameters, e.g. *L* or *τ*, as for the straight cable. Note that this resonance is similar to what we found in the oblique dendrites of the pyramidal cell.

Besides the bending angle Θ, the presence or not of a resonance is solely determined by the length of the bent branch, *D*, relative to the length of the main branch, *H* (Fig. 3 E,F and Fig. S5). The resonance appears when the sensitivity to low frequency fields decreases. The higher the resonance frequency, the lower the resonance peak. However, the amplitude of the resonance amplitude, i.e. the peak minus the sensitivity to low frequency fields, depends both on the resonance frequency and on the decrease of sensitivity to DC fields. While the membrane time constant *τ* determines the resonance frequency, it has no effect on the presence of a resonance and a very limited impact on its amplitude.

To get a better insight in the mechanism behind this resonance, we look at the field sensitivity distribution along the bent cable (Fig. 3D). For a given frequency, the field sensitivity reaches a local minimum on both branches. For high frequency fields, i.e. low space constant λ_*gen*_(*ƒ*_*t*_), there is little interaction between both branches. The local minima are therefore located closer to the middle of the branches, as in the straight cable case. On the contrary, for low frequency fields, i.e. high λ_*gen*_(*ƒ*_*t*_), the branches exert a stronger influence on each other, which shifts the locations of their respective field frequency local minima. Reducing the membrane time constant lowers the effects of the frequency on λ_*gen*_(*ƒ*_*t*_) such that both branches interact with each other up to higher frequencies (Fig. S6).

In summary, we showed that in absence of active channels, the apical dendrites of a pyramidal cell tends to be more sensitive to low frequency fields oriented parallel to the somato-dendritic axis than the basal dendrites and the soma. This difference, which disappears for higher frequencies, can be explained by the presence of radial branches close to the soma. We also found a “passive resonance” in some oblique and basal dendrites that have an acute branching angle with the apical axis. We explain this resonance as a competition between the main branch (i.e. the apical dendrites) and the bent branches (oblique and basal dendrites) that experience the fields in opposite directions.

### Role of active properties

#### Biophysical model with experimentally constrained conductances

After investigating the role of the morphology of a passive pyramidal neuron in its sensitivity to weak electric fields, we now analyze the effects of active membrane properties. We consider the original *Hay et al.* [16] model, i.e the same reconstructed morphology as above including active conductances that were constrained using experimental data. Similarly to the passive case, we investigate the subthreshold sensitivity of the active cell, i.e. with all active channels, to an electric field. In absence of synaptic input or extracellular field, the membrane in the apical dendrite of this model is more depolarized than in the basal dendrites and the soma (see Fig. S8A in Supporting information). Fig. 4A displays the effective membrane polarization due to a DC field, i.e. the membrane polarization around the rest value of each location. Unlike in the passive model, the response of the apical dendrites to DC fields is no longer of higher amplitude than at the soma and basal dendrites. Moreover the membrane potential presents a strong overshoot at the field onset in the apical dendrite.

**Figure 4.**
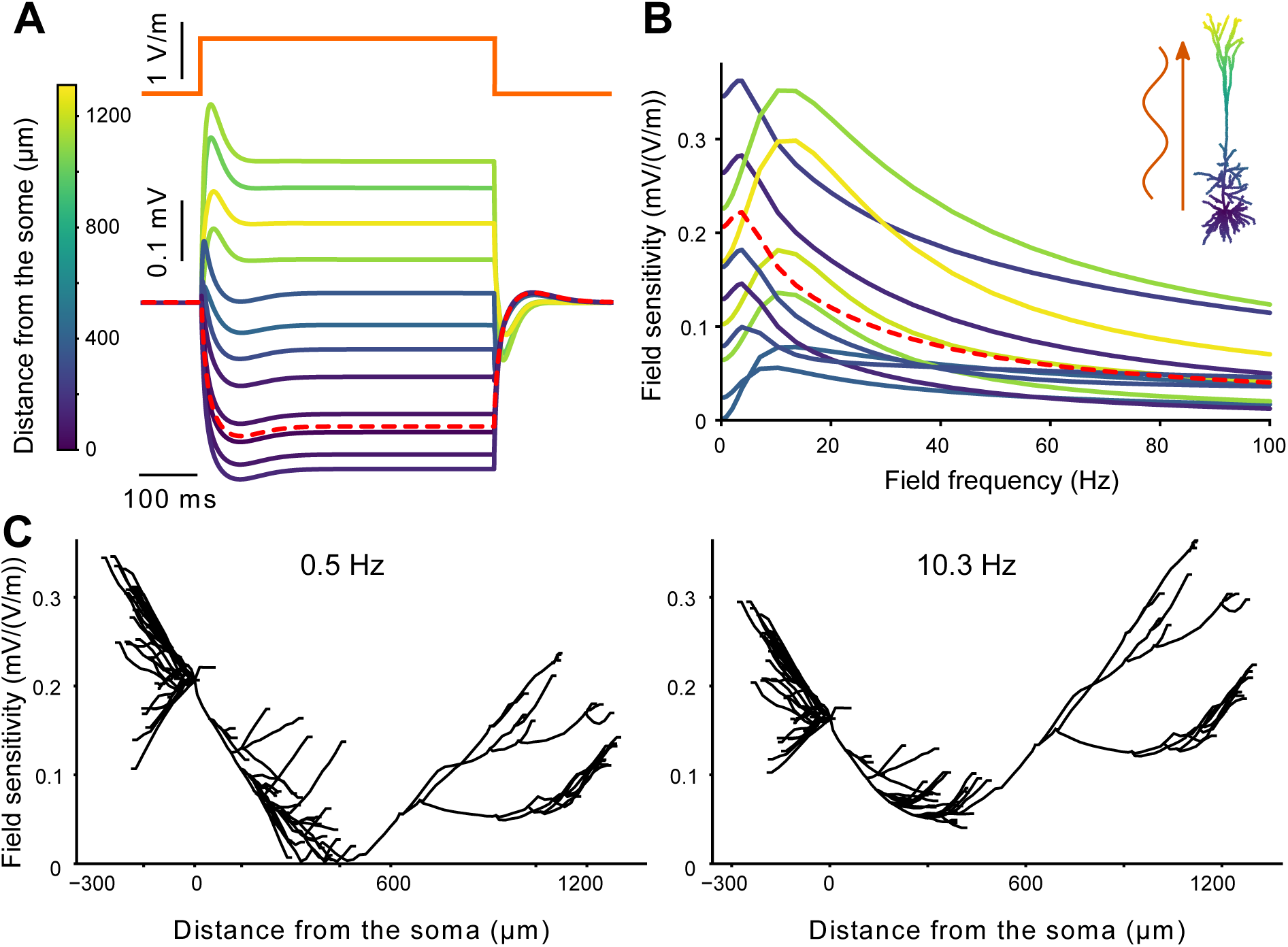
Unlike the soma and basal dendrites, the apical dendrites of an active cell have an increased subthreshold response to AC field of 10-20Hz. We consider a pyramidal cell model of Hay et al. [16] with all active channels. (A) Membrane polarization around the resting state due to a positive step current electric field (orange). (B) Frequency-dependent sensitivity of the cell to AC fields measured at different location on the whole cell. (C) Sensitivity to sinusoidal fields of frequency 0.5Hz (left) and 10.3Hz (right) as a function of distance to the soma. For clarity basal dendrites are plotted with negative distance. Colors of the polarization (A) and field sensitivity (B) correspond to the distance from the soma as depicted in A. The red dashed lines correspond to the soma.

Compared to the passive case, the sensitivity of the apical dendrite to low frequency AC fields is decreased (Fig. 4B). This produces a strong frequency resonance in the apical field sensitivity in the 10-20 Hz range. Basal dendrites and soma do not present this strong resonance but have a slight resonance around 5 Hz. While the basal dendrites and the soma are more sensitive to low frequency fields than the apical dendrites (Fig. 4C,D), the opposite is true at the resonance frequency of the apical dendrites when basal dendrites and soma are less sensitive than the apical dendrites. Locations which already presented a frequency resonance in the passive case, namely some proximal dendrites, still display qualitatively the same field sensitivity profile in the active model.

Fig. 5 shows a comparison of the frequency-dependent field sensitivity of the passive and active models at a selected subset of locations. To get a better understanding of the role of active channels on the cell field sensitivity profile we consider the impact of the three main differences between the active and passive models: (i) the non-uniformity of the membrane potential at rest, (ii) the increased resting conductance resulting from the additional active channels and (iii) the dynamics of the active channel themselves.

**Figure 5.**
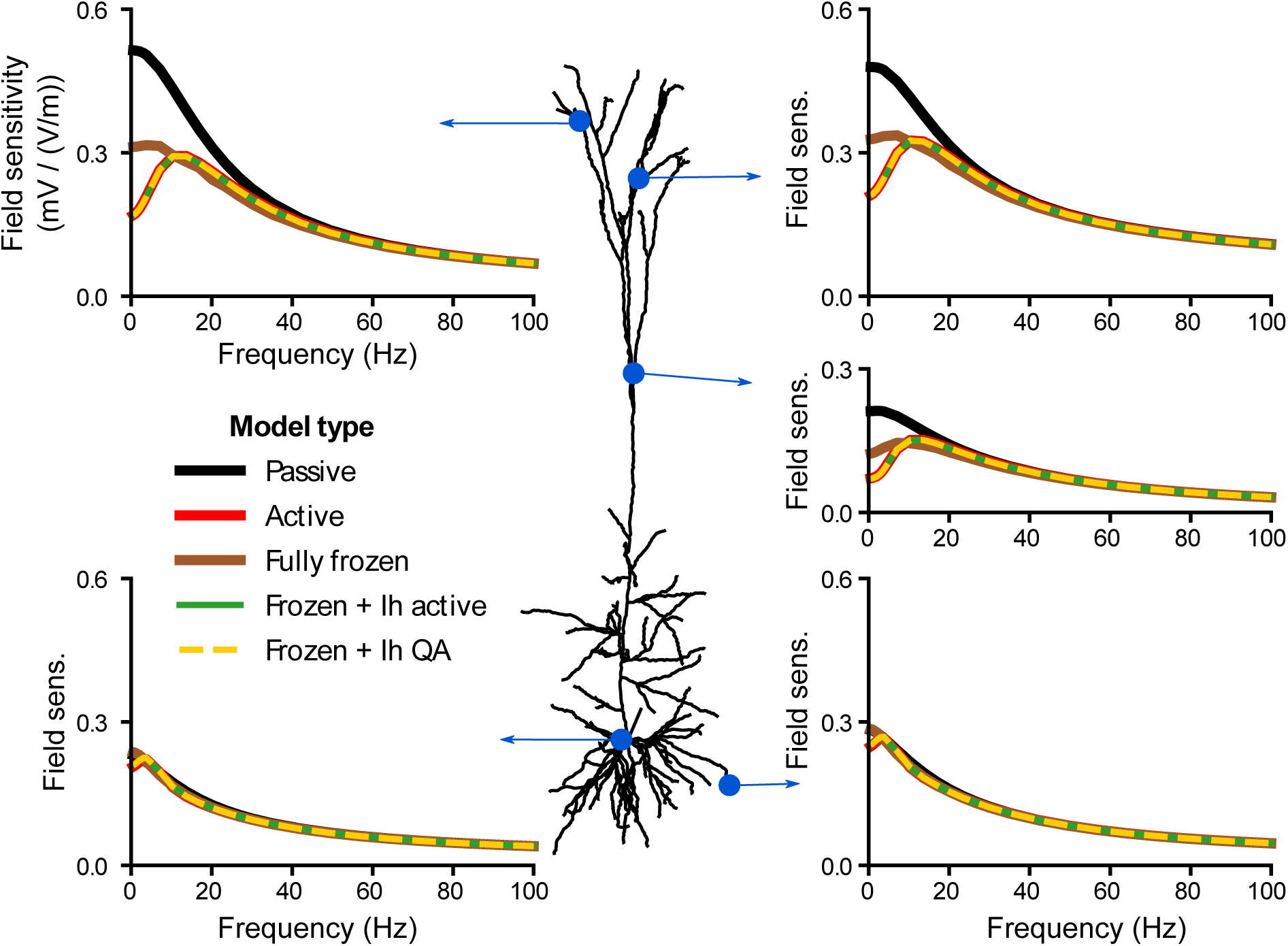
The hyperpolarization-activated inward current, *I*_*h*_, is responsible for the frequency resonance in the sensitivity of apical dendrites to AC fields. The subplots display the field sensitivity (in mV/(V/m)) of the cell at given locations depending on the channels included in the model: without any active channels (*passive*) or with all active channels present in the model of Hay et al. [16] active (*Active*). We also consider the model with frozen channels, i.e. their gating variable is fixed to their resting value. We freeze either all the channels (*Fully frozen*) or all except the *I*_*h*_ which is fully active (*Frozen* + *I*_*h*_ *active*) or linearized following the quasi-active approximation (*Frozen* + *I*_*h*_ *QA*).

Adjusting the leak reversal potential along the membrane, we uniformly set the membrane potential to various values. The field sensitivities at the apical tuft of these models do not differ qualitatively from the one of the original model: they all display a resonance in the apical tuft (see Fig. S8 in Supporting Information), though the frequency and amplitude of the resonance vary with the resting membrane potential. For highly depolarized resting membrane potential, the somatic sensitivity to low frequency fields drastically increased.

The next difference between the active and passive models resides in the increased resting conductance due to the presence of active ion channels. To evaluate its impact on the cell sensitivity to AC fields, we freeze the active channels, i.e. fix their gating variables to their value at rest [19] (see Methods). Frozen channels simply act as additional leak conductances. Compared to the passive model (Fig. 5, black solid curve), the addition of frozen channels (brown curve) decreases the sensitivity of the apical dendrites to fields with frequencies up to 40 Hz. At these locations, the field sensitivity of the fully frozen model is halfway between that of the passive and active model (red curve). Despite presenting the same field sensitivity as the active model for frequencies higher than the resonance, the frozen model still does not display any strong resonance. The field sensitivity at the soma and basal dendrites is little affected by the addition of frozen channels.

We further consider the fully frozen model and reactivate, i.e. unfreeze, each channel separately in order to test the role of single channels active properties on the resonance. Unfreezing the hyperpolarization-activated inward current *I*_*h*_ turns out to be sufficient to obtain the resonance and, in fact, fully accounts for the field sensitivity of the active model (Fig. 5, green curve). Unfreezing other channels and keeping *I*_*h*_ frozen does not yield any resonance.

In summary, in the absence of active channels the field sensitivity of the neuron model mostly decreases with frequency. Introducing the active conductances reduces the sensitivity of the apical dendrites to low frequency fields and produces a resonance. We attribute this to the increased resting conductance provided by the active channels and to the dynamics of the *I*_*h*_ channel. The exact amplitude and position of the resonance depends on the membrane potential at rest.

### Quasi-active case

The results obtained in the preceding section strongly depend on the membrane channels present in the model by Hay et al. [16]. In order to generalize these results to any kind of subthreshold current, we consider a simpler channel model and conduct a systematic study of its effects on the model field sensitivity. Specifically, we use the quasi-active (QA) approximation [19–21], which is a linearization approximation of active currents around the rest membrane potential.

Let’s consider a neuron model with a leak current and a single voltage-dependent channel. The dynamics of the channel’s gating variable and membrane potential can be linearized around the rest value *V*_*R*_ (see Methods). The resulting membrane potential of a given cell compartment is then governed by the
equation:

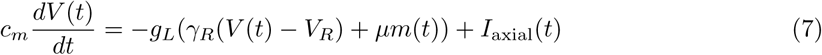

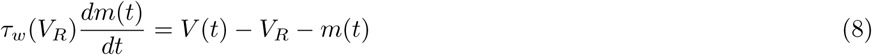

where *I*_axial_ is the axial current coming from neighboring compartments, *c*_*m*_ the membrane capacitance and *g*_*L*_ the leak conductance. The parameters of the quasi-active current, namely *γ*_*R*_, *μ* and *τ*_*w*_(*V*_*R*_), are independent on the membrane potential and fully determined by the active channel parameters at *V*_*R*_. Before going further, we assess the validity of the linearization approximation for the present study. We compute the field sensitivity of the preceding model with all channels frozen except the *I*_*h*_ channel, which we linearize using the QA approximation. The linearized model perfectly replicates the resonance in the apical dendrite (Fig. 5, yellow dashed curve). Due to the weakness of the considered field and consequently of the resulting membrane polarization, the active channels indeed operate in a near linear regime, hence, the QA approximation is valid for our case.

A key advantage of the QA framework is that it provides a strong reduction of the number of parameters. For low frequency stimulations, the sign of *μ* fully determines the behavior of the channel [19, 22]. If *μ* is negative, the channel provides positive feedback to modulations of the membrane; the current is *regenerative*. On the contrary if *μ* is positive, the channel provides negative feedback to membrane modulations and the current is *restorative*. In case *μ* equals zero, the dynamics of the normalized gating variable m has no influence on the membrane; the quasi-active current acts as a passive leak current (i.e., identical to a frozen active current). Three factors can influence the effects of QA channels on the cell field sensitivity: the spatial distribution of the QA conductances, the channel type (i.e. the sign of *μ*), and the channel activation time constant *τ*_*w*_(*V*_*R*_). We first consider a passive QA channel (*μ* = 0) with three different conductance distributions: 1) uniform over the whole cell, 2) linearly increasing with distance from the soma, and 3) linearly decreasing with distance from the soma (see Methods). We parametrize all 3 distributions to have the same sum of conductance over the entire neuron model. The conductance distribution mainly affects the sensitivity to fields below 80Hz (Fig. 6). In general, the higher the local conductance, the lower is the field sensitivity. Indeed, the linearly increasing distribution of the passive QA channel results in a decreased field sensitivity at the apical dendrites and an increased field sensitivity at the soma and basal dendrites. The linearly decreasing conductance distribution have the opposite effect.

**Figure 6.**
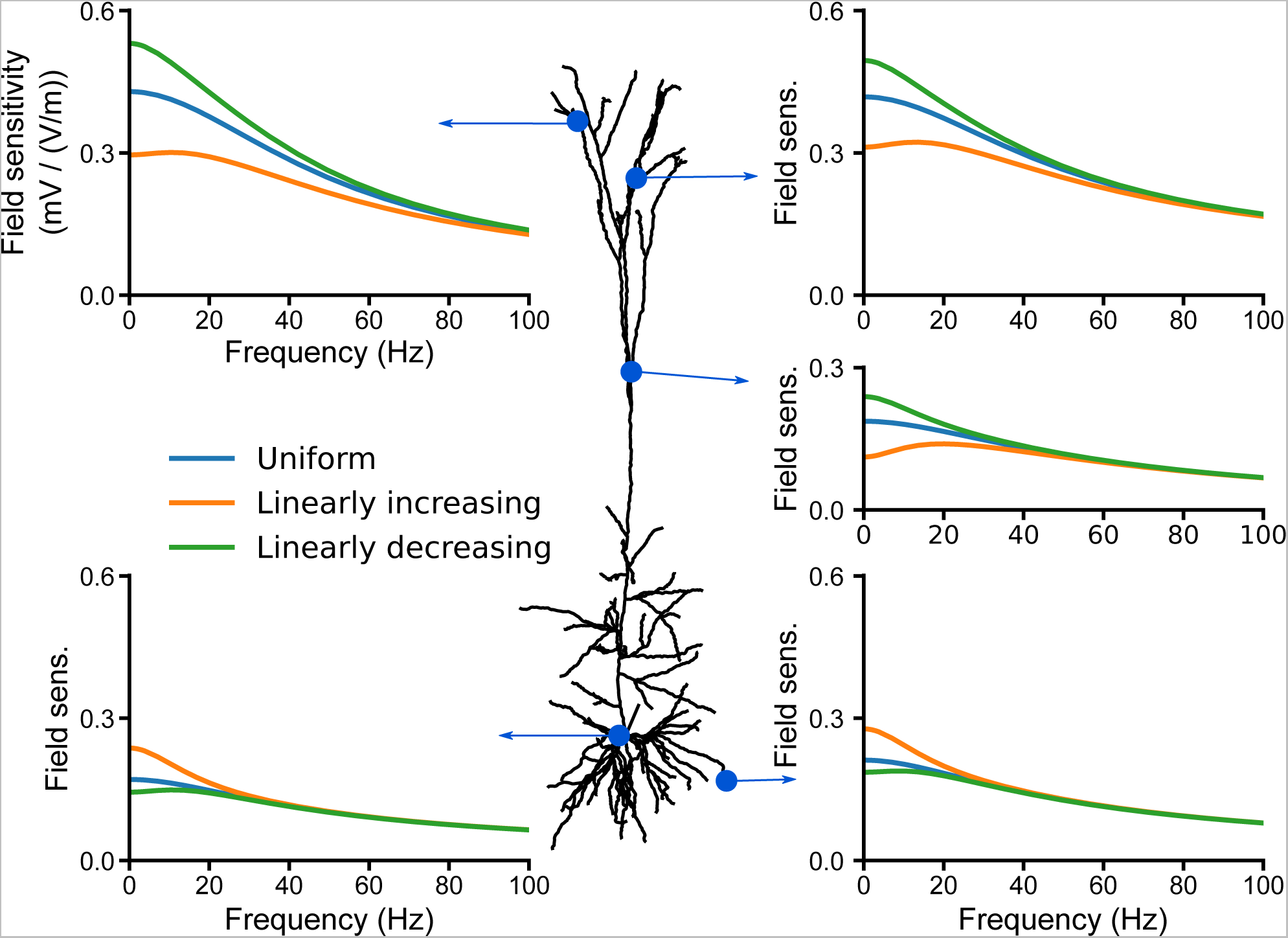
The quasi-active channel conductance distribution affects the cell field sensitivity depending of the local conductance at the considered location. We consider a neuron model which includes solely a leak conductance and a single quasi-active channel (QA). The QA channel have no active dynamics (*μ* = 0) and acts as an additional leak current. We consider 3 different QA conductance distributions: uniform and linearly increasing/decreasing with distances from the soma. For each distribution, the sum over the whole cell of the QA conductances at rest is equal to the sum of the leak conductances. The subplots display the cell field sensitivity (in mV/(V/m)) for the different conductance distributions.

We investigate the impact of the channel type on the field sensitivity. We next consider a uniformly distributed QA channel with different values of *μ*. Due to their negative feedback to modulations, restorative currents (*μ* > 0) reduce the sensitivity to low frequency fields compared to passive channels (Fig. 7). This decrease induces a frequency resonance in the field sensitivity profile similar to what we observed with *I*_*h*_, which is indeed a restorative current. In contrast, the positive feedback of regenerative current increases the cell sensitivity to low frequency fields, hence, no resonance appears. Due to the uniform distribution of the channel conductance, we observe this effect at all considered points on the neuron (soma, basal and apical dendrite).

**Figure 7.**
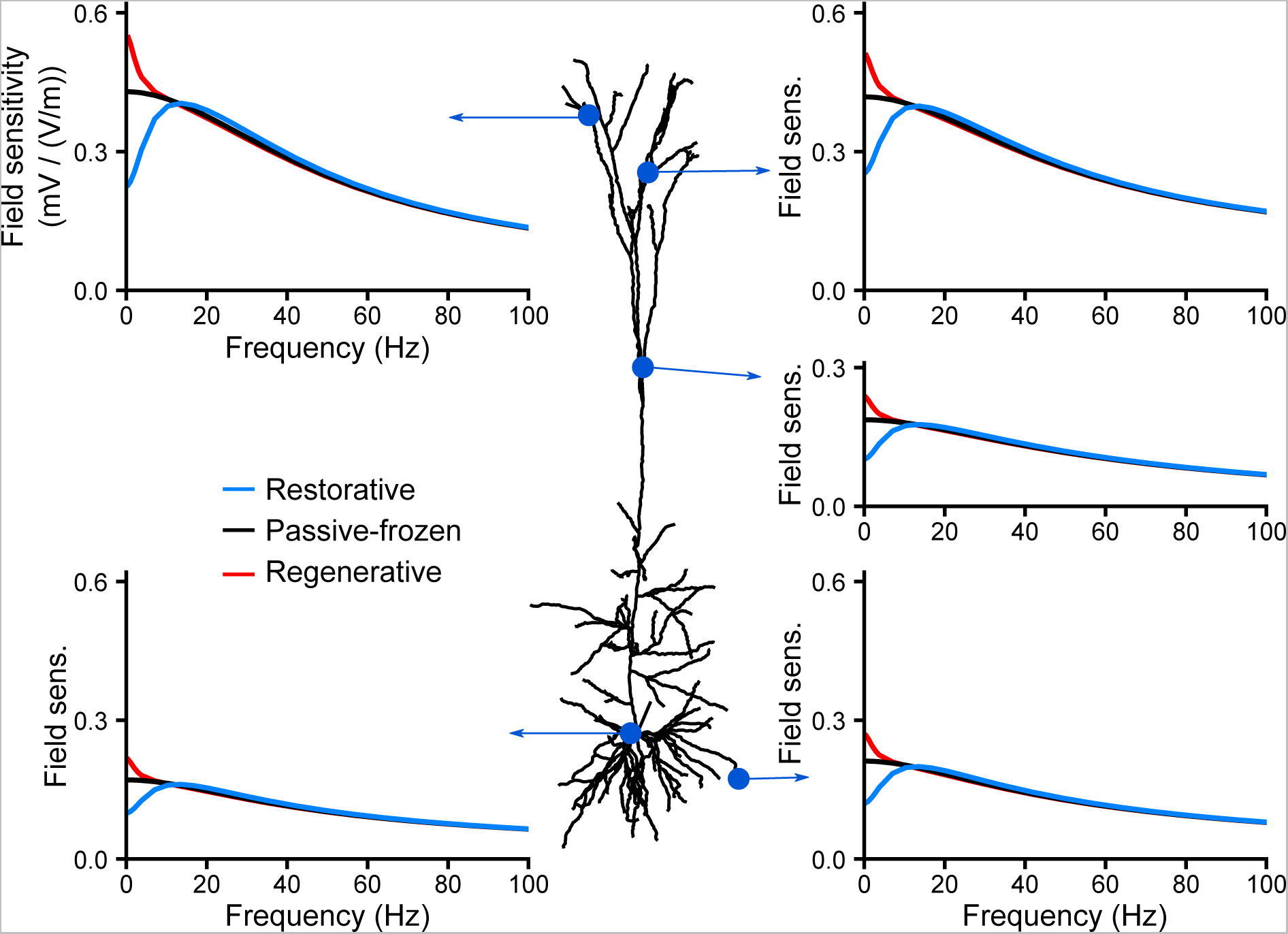
While regenerative currents increase the cell sensitivity to low frequency fields, restorative currents decrease it and induce a resonance. We consider a neuron model which includes solely a leak conductance and a single uniformly distributed quasi-active channel (QA). The plots display the cell’s sensitivity (in mV/(V/m)) to AC fields at different location in case of restorative (blue, *μ*\* = 2), passive (black, *μ*\* = 0) and regenerative (red, *μ*\* = −1) quasi-active currents.

The observed effects depend on the distribution of channels across the neuron. For a linearly increasing distribution, this change of the field sensitivity is mostly observed in the apical tree, while the basal and somatic compartments are hardly affected by the channel dynamics (Fig. S9 in Supporting Information). In contrast, for a linearly decreasing distribution, the field sensitivity at the soma and basal dendrites vary with the channel dynamics (Fig. S10 in Supporting Information), while it is less affected at the apical dendrites compared to the linearly increasing conductance distribution case. Overall, these results highlight the local effects of the QA channels: they affect the field sensitivity most strongly at locations where the channels have the highest density.

We finally look at the effect of the channel activation time constant *τ*_*w*_(*V*_*R*_) on the field sensitivity profile 8. Faster channels (lower values of *τ*_*w*_(*V*_*R*_)) are able to modulate the field sensitivity up to higher field frequencies. Restorative currents also act as a low-pass filter whose cut-off frequency is determined by the activation time constant. It directly affects the resonance frequency: the faster the activation time constant *τ*_*w*_(*V*_*R*_), the higher the resonance frequency. In the case of a fast channel time constant, e.g. *τ*_*w*_(*V*_*R*_) = 1 ms, the resonance disappears because the active current dynamics attenuate the field sensitivity over the entire frequency range.

**Figure 8.**
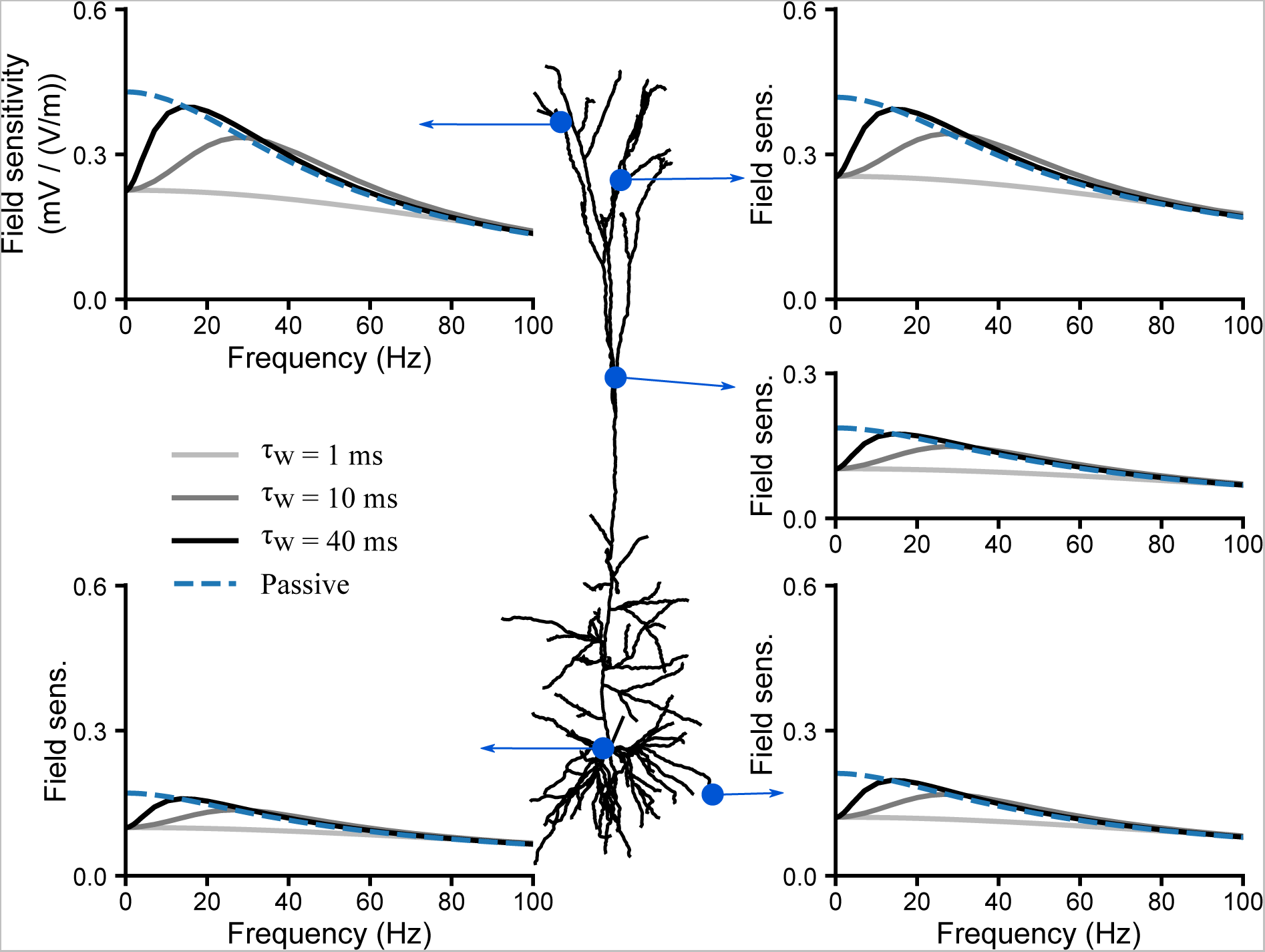
The time constant of restorative currents determines the resonance frequency of the sensitivity to AC fields. We consider a neuron model which includes solely a leak conductance and a single uniformly distributed quasi-active channel (QA). The plots display the cell’s sensitivity (in mV/(V/m)) to AC fields at different locations in case of a passive (*μ*\* = 0, dashed line) or a restorative (*μ*\* = 2, dashed lines) quasi-active current. The shades of grey encode the restorative QA channel time constant, *τ*_*w*_(*V*_*R*_).

## Discussion

In the present modelling study, we investigated the subthreshold frequency-dependent sensitivity of pyramidal neurons to weak extracellular electric fields. We found a strong frequency resonance around 10-20 Hz in the field sensitivity at the apical dendrites, which is absent at the soma and basal dendrites. We attributed this differential frequency-dependence of the field sensitivity to a concurrent (i) increase of field sensitivity at the apical dendrite due to the morphology of pyramidal cells and (ii) a decrease of this sensitivity to low frequency fields due to the presence of active channels, specifically, the h-type conductance. Our systematic analysis of the mechanism behind the observed resonance provides a better understanding of the relative role of morphology and membrane properties on the sensitivity of cortical neurons to weak electric fields. Below, we further summarize our findings and relate them to previous studies.

### Impact of the morphology

In accordance with previous studies [8, 9] we found an opposite polarization in the apical dendrite compared to the soma and basal dendrites of a passive pyramidal cell model. At most locations, the passive cell’s sensitivity decreased monotonously with the field frequency. This suggests that the frequency-dependent field sensitivity profile measured in vitro at the soma [14] is a purely passive effect. Moreover, the cell sensitivity to low frequency fields parallel to the somato-dendritic axis is not symmetrical but is stronger in the apical dendrites. It results from the presence of numerous dendrites extending radially, e.g. oblique and basal dendrites, close to the soma. These branches reduce the field sensitivity close to their branching points as observed in the bent cable (Fig. S3 in Supporting information).

Interestingly, some basal and oblique branches showed a frequency resonance in their field sensitivity. We observed this resonance for branches that were pointing in the apical direction This resonance was of relatively small amplitude, hence, it is unlikely to greatly affect the cell response to tACS. Nevertheless, similar resonances could occur in spatially non-uniform fields such as endogeneous fields.

To explain the phenomena observed in the reconstructed cell, we considered a simpler model. Through application of Fourier transforms and the Green’s function (i.e., the impulse response function), we analytically derived the polarization of a passive cable due to time-dependent and spatially non-uniform fields (see also [23]). We first illustrated how the sensitivity of a straight cable depends on the field frequency and cable length (Fig. 2; see also [11,24]) and explained the findings using the frequency-dependent space constant, λ_*gen*_(*ƒ*_*t*_) (Eq. 5). This generalized space constant is also useful to explain the frequency resonance observed in the passive cable with an acute bending angle (Fig. 3). In this case, the bending induces a change in the field orientation experienced locally by the cable and reduces the sensitivity to DC fields while not affecting sensitivity to alternating fields. Similar effects could appear for spatially non-uniform fields, which could explain the ”passive resonance” observed in the sensitivity of a straight cable to a field generated by a point source [24].

### Impact of the active currents

We further investigated the polarization of different compartments of a pyramidal cell neuron with voltage-dependent ion channels. The sensitivity of this neuron model to AC and DC fields scaled linearly with the field amplitude, in agreement with in vitro recordings [9,14]. Also in accordance with in vitro measurements in CA1 pyramidal cells [9], our active model exhibited a stronger DC polarization at the basal dendrites and soma than in the apical dendrites. A strong overshoot was also present at the field onset in the apical dendrite. We found a slight resonance at 5 Hz in the somatic sensitivity to AC fields. This resonance can neither be confirmed nor excluded from available in vitro data (see Fig. 10 in [14]). Our model exhibited a strong resonance in the apical dendrites in the 10-20 Hz range. This resonance resulted from the presence of the *I*_*h*_ channel. There are, to our knowledge, no experimental studies that investigated the polarization of the dendritic tree due to AC fields.

We next considered a simplified model of voltage-dependent ion channels, the quasi-active approximation [20, 21] to systematically investigate the effects of active channels on the sensitivity to electric fields. This linear approximation is particularly applicable for our purposes, regarding the low amplitude of the field produced during transcranial current stimulation, which results in small polarizations of the cell membrane. The quasi-active approximation emphasizes that there are two classes of voltage-dependent currents: regenerative and restorative currents. The restorative currents oppose slow voltage fluctuations and consequently tend to decrease the sensitivity to low frequency fields. This results in a frequency resonance as observed with the *I*_*h*_ channel, which is indeed a restorative current. The presence of a regenerative current (e.g., a persistent sodium current) induces the opposite effects, amplifying the polarization in response to low-frequency fields, therefore not leading to resonances. These effects of the channel type also apply to other pyramidal cell morphologies (Fig. S11 and Fig. S12 in Supporting Information).

The amplitude of the field sensitivity decrease or increase depended on the slope of the channel activation function at rest and the density of the active current (relative to the other currents). The channel activation time constant at rest determines the cutoff frequency. This explained the variation of the resonance frequency and amplitude in the model by [16] when changing the resting membrane potential (Fig. S8 in Supporting Information).

### Effect of the polarization on spiking activity

In this work, we focused exclusively on the sub-threshold response of neurons to extracellular fields. The strong frequency resonance we highlighted was exclusively present in the apical dendrite. Whether this resonance can modulate the output spiking activity still needs to be addressed. Following the ”somatic doctrine” [7], most of the modelling and experimental studies have solely considered the effects of a somatic polarization on neural output activity (see [25] for a review). Field-induced somatic polarization has been shown to affect spike-timing as a function of the field strength for low-frequency fields, and to induce coherent firing for high frequency fields [10]. Recently, the polarization of dendritic endings were found to modulate synaptic efficacy in vitro [13]. Though the observed effect was very subtle (±1.17% of EPSP amplitude change per V.m^−1^), this modulation could be enhanced at the network scale, each neuron being subject to the same field. Similarly, DC fields have been shown to affect plasticity by modulating long term potentiation (LTP) and depression (LTD) at Schaffer collaterals in CA1 rat hippocampal slices [12]. The nature of the modulation, i.e. enhancement or reduction, depended on the field orientation and the synapse location. These studies only considered DC fields and, in light of the present results, a stronger modulation could potentially be achieved using AC fields at the resonance frequency.

A potential effect of the dendritic polarization could also be the modulation of NMDA spikes, calcium spikes, and sodium spikes in the dendritic arbor [15, 26]. These events have a voltage threshold and greatly enhance the importance of distal dendritic input compared to proximal input. In particular, calcium spikes, which are generated exclusively in the apical dendrites, play a great role in the integration of feedback input from higher level areas in the cortex [27]. The generation of these spikes could be affected by the subthreshold resonance we found in the apical dendrite. In this case, through adjustment of the stimulation frequency, one could act on the relative strength of the stimulation at the apical and basal dendrite: choosing DC or very low frequency field, one would preferentially polarize the basal dendrites and the soma, while stimulation in the 10-20 Hz range would polarize the apical dendrites more strongly (cf Fig. 4C for the subthreshold response). Hence, this would allow to differentially polarize proximal dendrites versus distal apical dendrites, thereby modulating the balance between feedforward and feedback synaptic input [27].

Note however, that the translation of subthreshold resonance into a suprathreshold resonance is not trivial. For example, in a simple ball-and-stick model, the suprathreshold modulation was found to display a resonance which was not present in the subthreshold sensitivity [11].

## Conclusion

In conclusion, our modelling work provides an improved understanding of the role of neuron morphology and active and passive membrane properties in the subthreshold sensitivity of pyramidal neurons to weak electric fields. We showed that dendrites present a differential frequency-dependent polarization in response to fluctuating electric fields. Most importantly, our prediction of a strong electric field resonance in the apical dendrites could open the door to an improved design of tACS stimulation.

## Methods

### Complex cell model

#### Active cell model

We simulate the membrane polarization of a complex pyramidal cell neuron model using NEURON [28]. We consider the model published by Hay *et al.* [16]. This state-of-the-art model of a thick-tufted layer 5b pyramidal cell contains 9 ionic channels. The distribution of their conductances, except for *I*_*h*_, was fitted using multi-objective evolutionary algorithm to reproduce perisomatic response to current step input and back-propagating action potential-activated calcium spikes as reported experimentally. By default, the active cell model has a non-uniform rest membrane potential. When notified in the text, we adjust the leak reversal potential to obtain a uniformly distributed rest membrane potential at an arbitrary value [28]. For part of the analysis, we block the dynamics of given active channels, we refer to these channels as ”frozen”. The ”frozen” channels have their conductances and gating variables fixed to their rest values and therefore act as additional leak currents.

#### Quasi-active approximation

In order to lower the parameter state of active ion channels, it is possible to linearize them around the rest value, while still keeping their active properties [19,20,22]. This quasi-active approximation is here justified by the low amplitude of the considered electric fields and their small perturbations of the membrane potential.

Let’s consider a neuron with a leak current and a single active conductance. The membrane dynamics of a single segment of the cell can be described as:

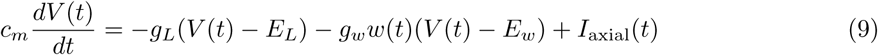

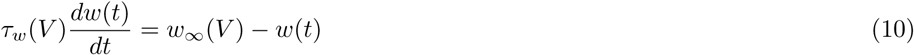

where *V*(*t*) = *V*_*i*_(*t*) – *V*_*e*_(*t*) is the membrane potential, *c*_*m*_ the membrane capacitance. *g*_*α*_, *E*_*α*_ (with *α* ∈ {*L, w*}) are respectively the conductances and reversal potentials of the leak (subscript *L*) and active (subscript *w*) currents. *I*_axial_ represents axial currents coming from neighboring segments. Here *g*_*w*_ is static and the dynamics of the active current are contained in the gating variable *w*, with time constant *τ*_*w*_(*V*) and activation function *w*_∞_(*V*).

We now linearize this equation using the Taylor expansion of *V* and *w* around the resting potential *V*_*R*_ and keeping only the first order term. Setting 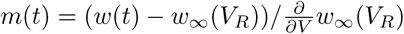, we obtain [19]:

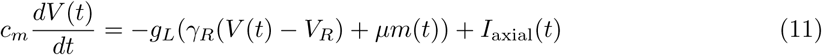

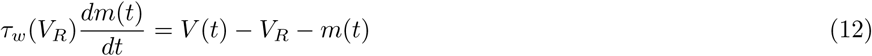

with 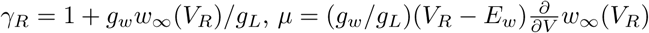.

This quasi-active approximation is still valid when several channels are present in the model. In this case each of them would be linearized separately.

To study the effects of the channel distribution on the neuron sensitivity, we consider several quasi-active channel distribution: (i) uniformly distributed, (ii) linearly increasing from the soma and (iii) linearly decreasing from the soma. We select the increasing and decreasing distributions to have a 60-fold difference between the soma and the most distal apical dendritic point [19]. This is in accordance with experimentally measured *I*_*h*_ distribution [29]. For all conductance distributions, we calibrate the total, i.e. summed over all cell compartments, quasi-active conductance *g*_*w*_ to be equal to the total leaky conductance *g*_*L*_. *g*_*L*_ is uniformly distributed and equal to 50*μS/cm*^2^. The resulting linearly increasing and decreasing distributions are defined respectively as *g*_*w*_(*x*) = 0.117*x* + 2.60 (*μS/cm*^2^) and *g*_*w*_(*x*) = −0.0539*x* + 71.9 (*μS/cm*^2^) where *x* (*μm*) is the distance along the dendritic tree between a segment and the soma.

Additionally, in order to define the type of the current *μ* independently of the local quasi-active channel conductance, we define *μ*^*\**^(*x*) = *μ*(*x*)*g*_*L*_/*g*_*w*_(*x*) such that 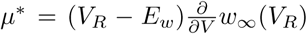. Unless specified otherwise, the quasi-active channel time constant is set to 50 ms. We also set the membrane capacitance *c*_*m*_ to 1 *μF/cm*^2^, the axial resistance *R*_axial_ = 100(Ω*cm*) and *w*_∞_(*V*_*R*_) = 0.5. V_R_ is set uniformly along the dendritic tree.

#### Simulation and computation of the sensitivity

We apply an extracellular field parallel to the somato-dendritic axis of the cell model. We set the extracellular potential *V*_*e*_(*x,t*) using the *extracellular* mechanism in NEURON. The somato-dendritic axis is determined by computing a volume-weighted vector between the soma and the cell center of mass. Similarly to the pasive cable case, we consider only spatially uniform DC and AC fields. For each field frequency, we first run the model for 700ms without the extracellular field in order to for the model to reach its steady state. A field of 1V/m was then applied for 2 cycles or at least 400ms and the last peak of polarization was used to determine the field sensitivity. Using longer simulations did not change the field sensitivity.

## Cable equation

### Membrane polarization due to an arbitrary field

We consider a cable of length *l* (mm) and radius *r* (mm). Its eletronic properties are characterized by the cable’s membrane time constant *τ* = *c*_*m*_/*g*_*m*_ (s) and electrotonic length scale 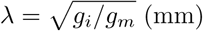, where *c*_*m*_ = 2*πrc* (F/mm) is the membrane capacitance per unit length and *g*_*i*_ = *ρ*_*i*_*πr*^2^(S · mm), *g*_*m*_ = 2*πrρ*_*m*_ (S/mm) are respectively the internal (axial) and membrane conductances per unit length. The parameters *c* (F · mm^2^), *ρ*_*i*_ (S/mm) and *ρ*_*m*_ (S/mm^2^) are respectively the specific membrane capacitance, the axial conductance and the membrane conductance. The cable membrane potential is defined as *v*(*x,t*) = *v*_*i*_(*x,t*) - *v*_*e*_(*x,t*). For simplicity we assume that the rest membrane potential is null.

The cable is subject to an electric field *E*(*x,t*) parallel to the cable axis, *x*:

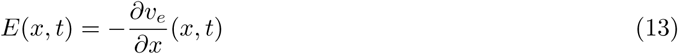

We do not assume any spatial non-uniformity or stationarity of the field but we do assume that the extracellular spatial and temporal components are independent:

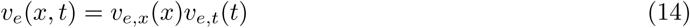

The dynamics of the membrane potential along the cable are governed by equation [17]:

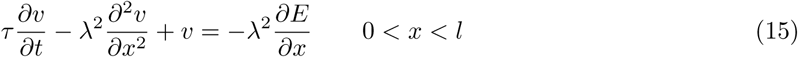

Rewritten in terms of *v*_*i*_ and *v*_*e*_:

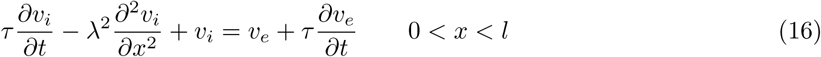

and the sealed end (no axial current flow) boundary conditions [23]:

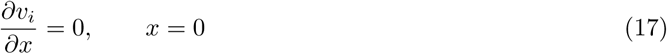

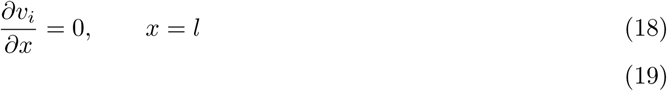

We solve the equation Eq. 16 using the separation of variables (for which we need the assumption of Eq. 14) and the temporal Fourier transform:

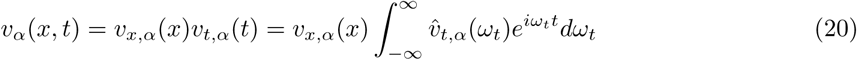

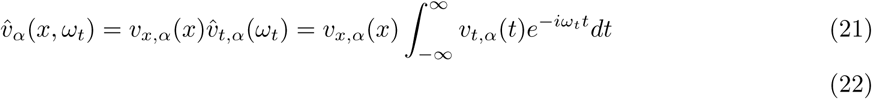

where *α* ∈ {*i, e*}, *ω*_*t*_ = 2*πƒ*_*t*_ denotes the temporal angular frequency and ^. the temporal fourier transform. Considering each angular frequency separately, i.e. for a fixed *ω*_*t*_, we rewrite Eq. 16 in the temporal Fourier domain:

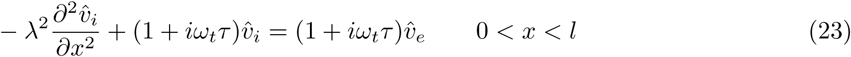

We define the complex *frequency-dependent space constant*, λ_*eq*_(*ƒ*_*t*_) as:

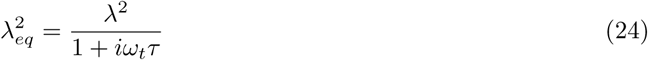

and the generalized frequency-dependent space constant:

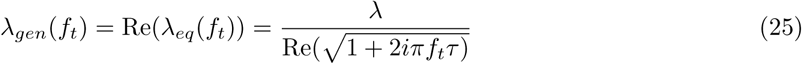

Eq. 23 becomes:

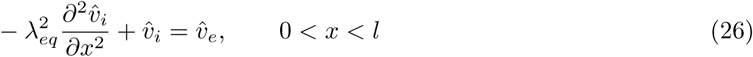

We now consider the spatial Fourier transform. To avoid confusion between imaginary terms coming from the temporal and spatial transforms, we use the cosine transform for the spatial domain:

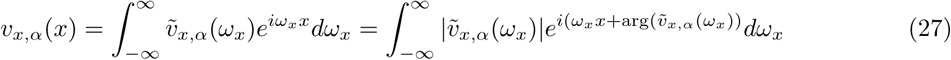

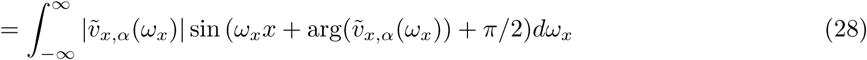

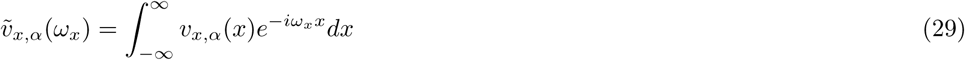

where *α* ∈ {*i, e*} and *ω*_*x*_ = 2*πƒ*_*x*_ denotes the spatial frequency of the electrical potential. This is valid since *v*_*x,α*_ is real, i.e. ∀*_x_*, *v*_*x,α*_(*x*) ∈ ℝ.

Taking each spatial component separately, i.e. for a fixed *ω*_*x*_, and setting 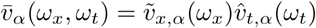 Eq. 26 becomes:

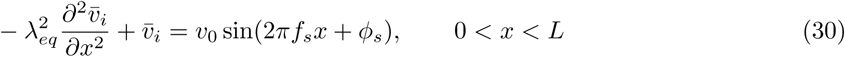

with 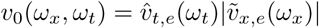 and 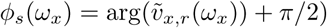. We then normalize the units:

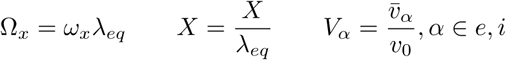

Eq. 30 becomes:

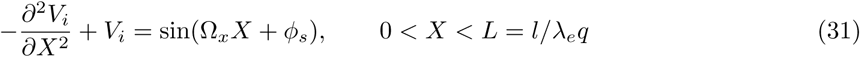

and the boundary conditions:

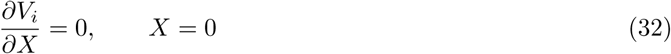

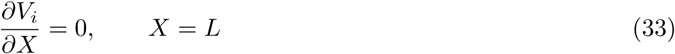

This equation can be solved using the Green’s function [23,30]:

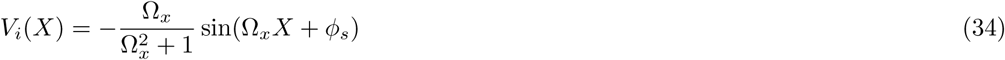

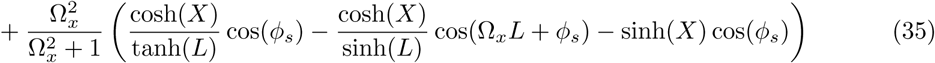

In terms of membrane potential *V* = *V*_*i*_ – *V*_*e*_

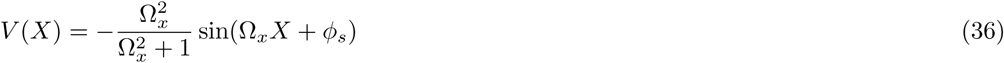

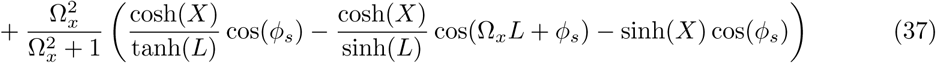

Here *V*(*X*) represent the subthreshold transmembrane potential variation along the cable due to an electric field. Its dependence in the field temporal, *ω*_*t*_, and spatial, *ω*_*x*_, angular frequencies are hidden in the variables *Ω*_*x*_, *φ*_*s*_ and λ_*eq*_. The full membrane modulation can be obtained by integrating over these angular frequencies.

The full equation for the membrane potential of the cable due to an arbitrary field with independent spatial and temporal component is:

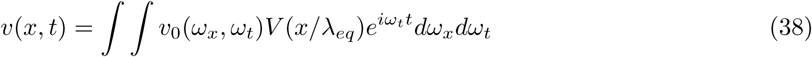

where *V* is a complex function given by Eq. 36 and it also depends on *ω*_*x*_.

The above solution solely requires the spatial and temporal components of the extracellular field to be independent. This assumption is not limiting the present work. Nevertheless, this restriction could be easily by-passed using the linearity of the cable equation and considering each spatial components separately.

**Field profiles** In this work, we consider DC or AC fields with fixed spatial profile

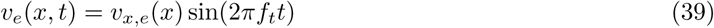
where *ƒ*_*t*_ ≥ 0. The resulting membrane polarization is of the form:

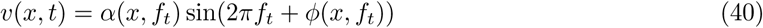

with:

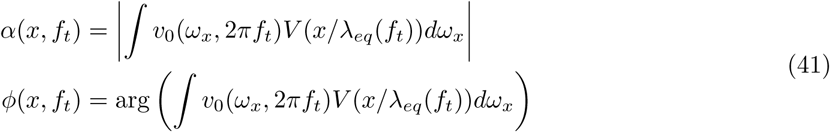

|*z*| and arg(*z*) correspond to the absolute value and argument of a complex number *z*. Moreover, we focus exclusively on spatially uniform extracellular field:

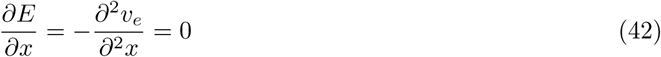

which means that for a straight cable, the extracellular potential increases or decreases linearly along the cable:

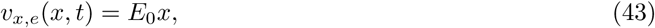

where *E*_*0*_ ∈ ℝ is the field amplitude in V/m. We also consider bent cables as sketched in Fig. 3. In this case the extracellular potential along the cable, can be obtained by projecting the bent part of the cable position on its axis:

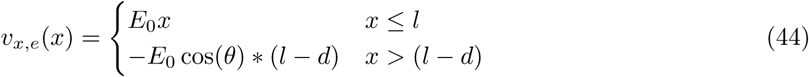
where *θ* is the bending angle in radian and *d* the length of the bent branch.

## Acknowledgments

This work was supported by the DFG Priority Programme SPP1665 (FA,KO), the Einstein Foundation Berlin (MR), and we want to thank Susanne Schreiber for her funding of MR through the Bundesministerium fr Bildung und Forschung (http://www.bmbf.de; grant number 01GQ0901 grant number 01GQ0901). The funders had no role in study design, data collection and analysis, decision to publish, or preparation of the manuscript.

---

**Figure S1.**
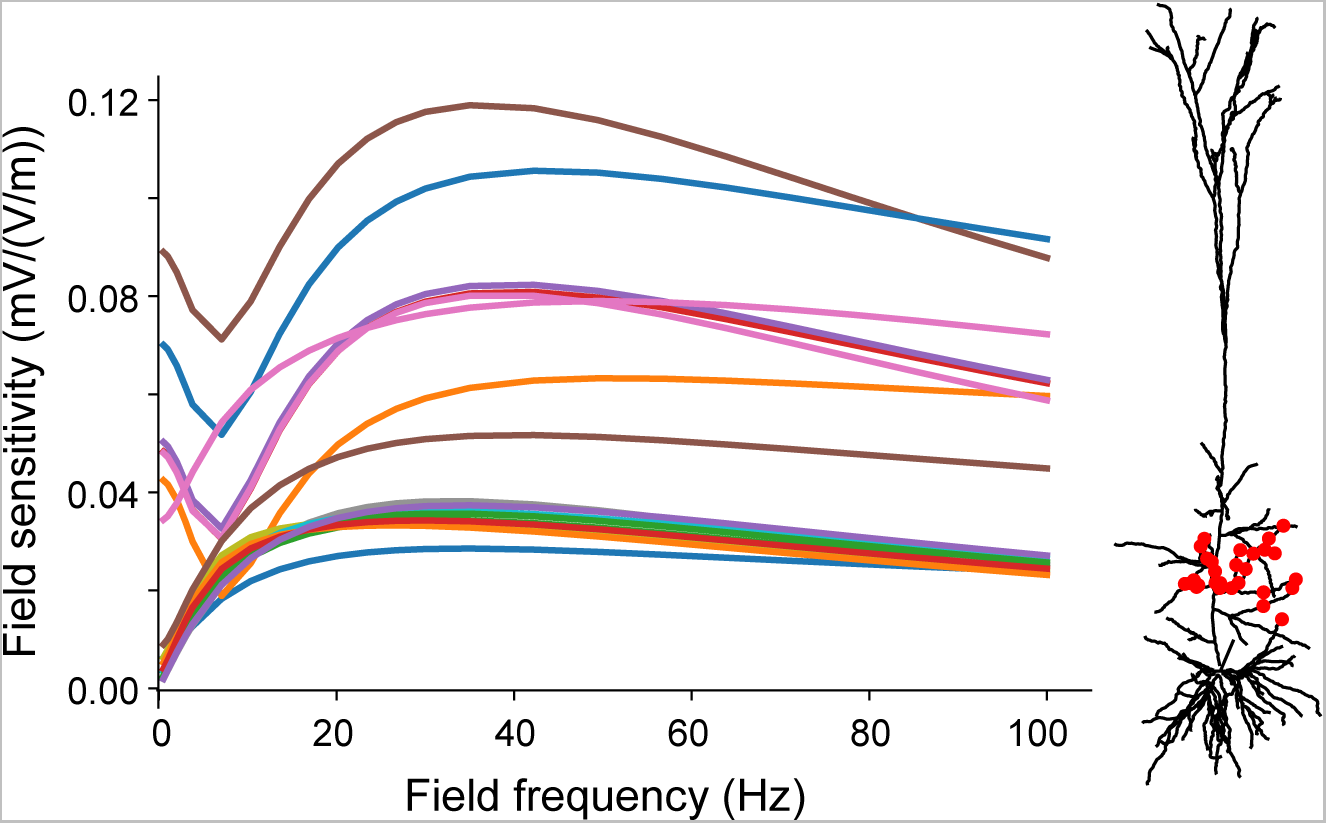
A passive pyramidal neuron exhibits a frequency resonance in its field sensitivity at the proximal dendrites. (Left) Frequency-dependent sensitivity of the passive cell, i.e. without any active ion channels, due to AC field parallel to the somato-dendritic axis. The locations where these sensitivities are measured are displayed on the cell (right).

**Figure S2.**
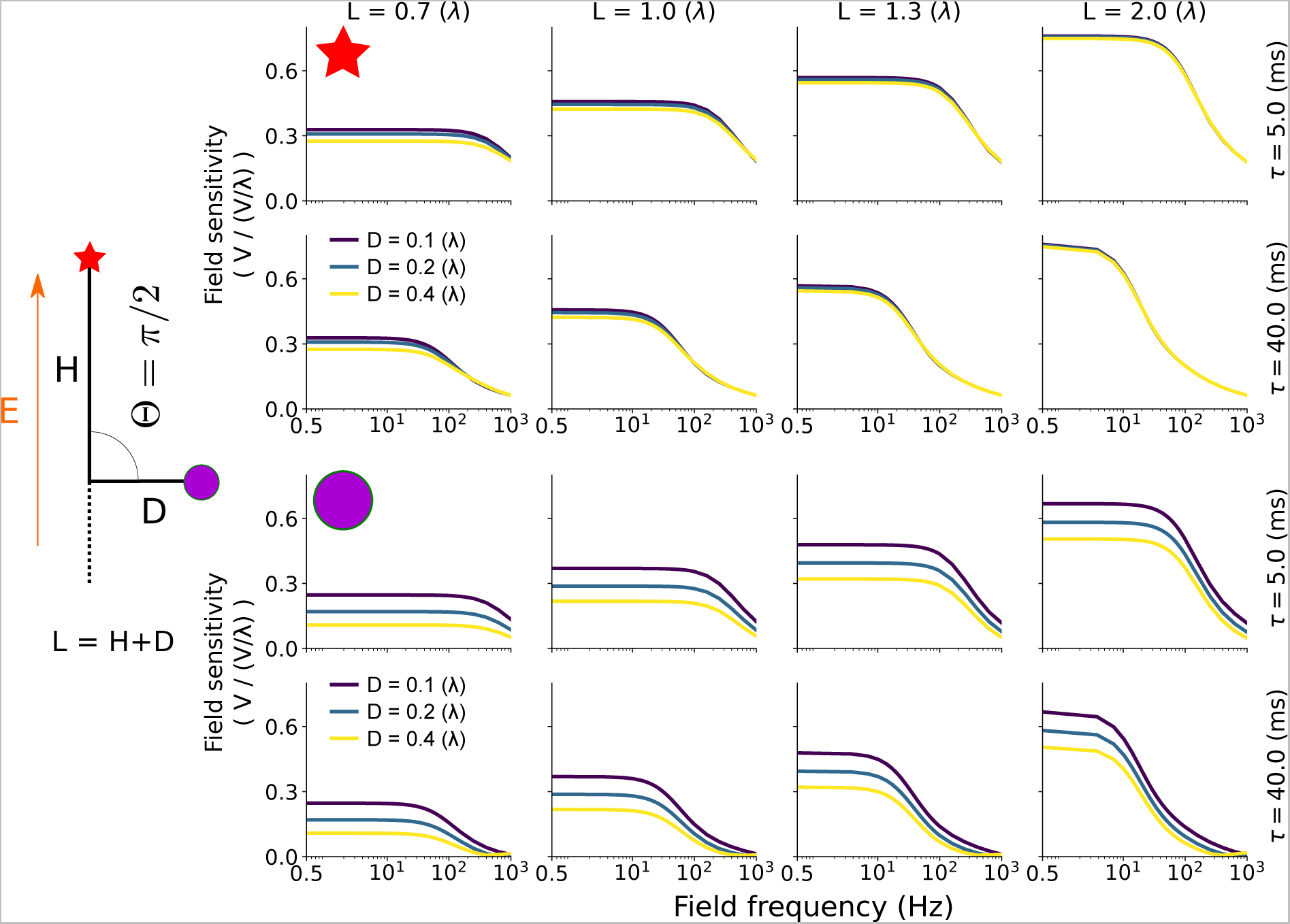
In a passive cable with a right bending angle, the length of the bent branch mainly affects the field sensitivity at its extremity, without inducing a resonance. The subplots represent the field sensitivity (in V / (V/λ)) at both cable ends: (top,red star) the unbent branch and (bottom, violet circle) the bent one, as function of the field frequency (x axis). The field sensitivity are displayed for various membrane time constant *τ* (rows for each location), total cable length *L* (increasing from left to right) and bent branch length *D* (color coded). The bending angle is Θ = *π*/2 (rad).

**Figure S3.**
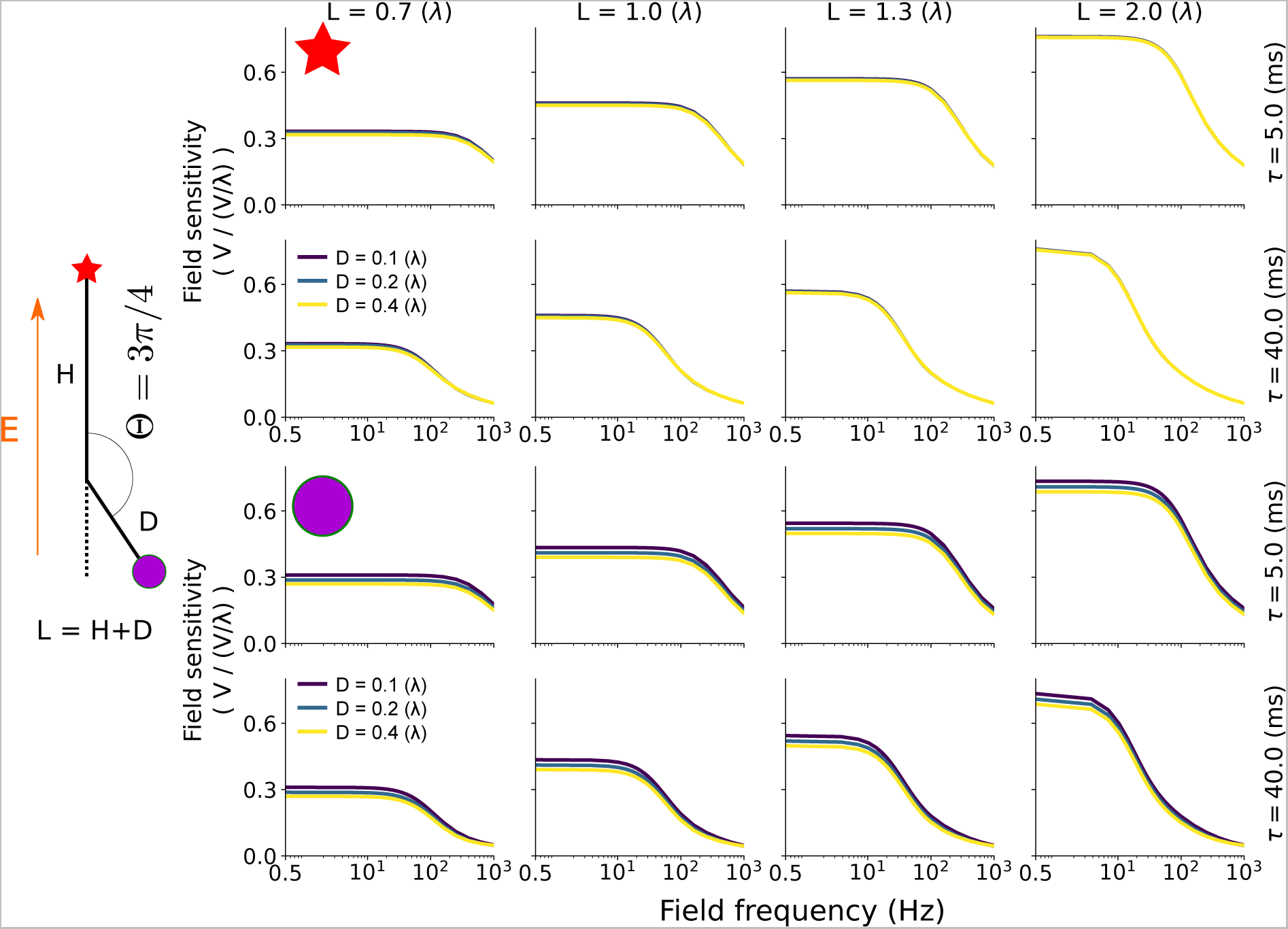
In a passive cable with an obtuse bending angle, the length of the bent branch has little effects on the field sensitivity at both extremities. The subplots represent the field sensitivity (in V / (V/λ)) at both cable ends: (top,red star) the unbent branch and (bottom, violet circle) the bent one, as function of the field frequency (x axis). The field sensitivity are displayed for various membrane time constant *τ* (rows for each location), total cable length *L* (increasing from left to right) and bent branch length *D* (color coded). The bending angle is Θ = 3*π*/4 (rad).

**Figure S4.**
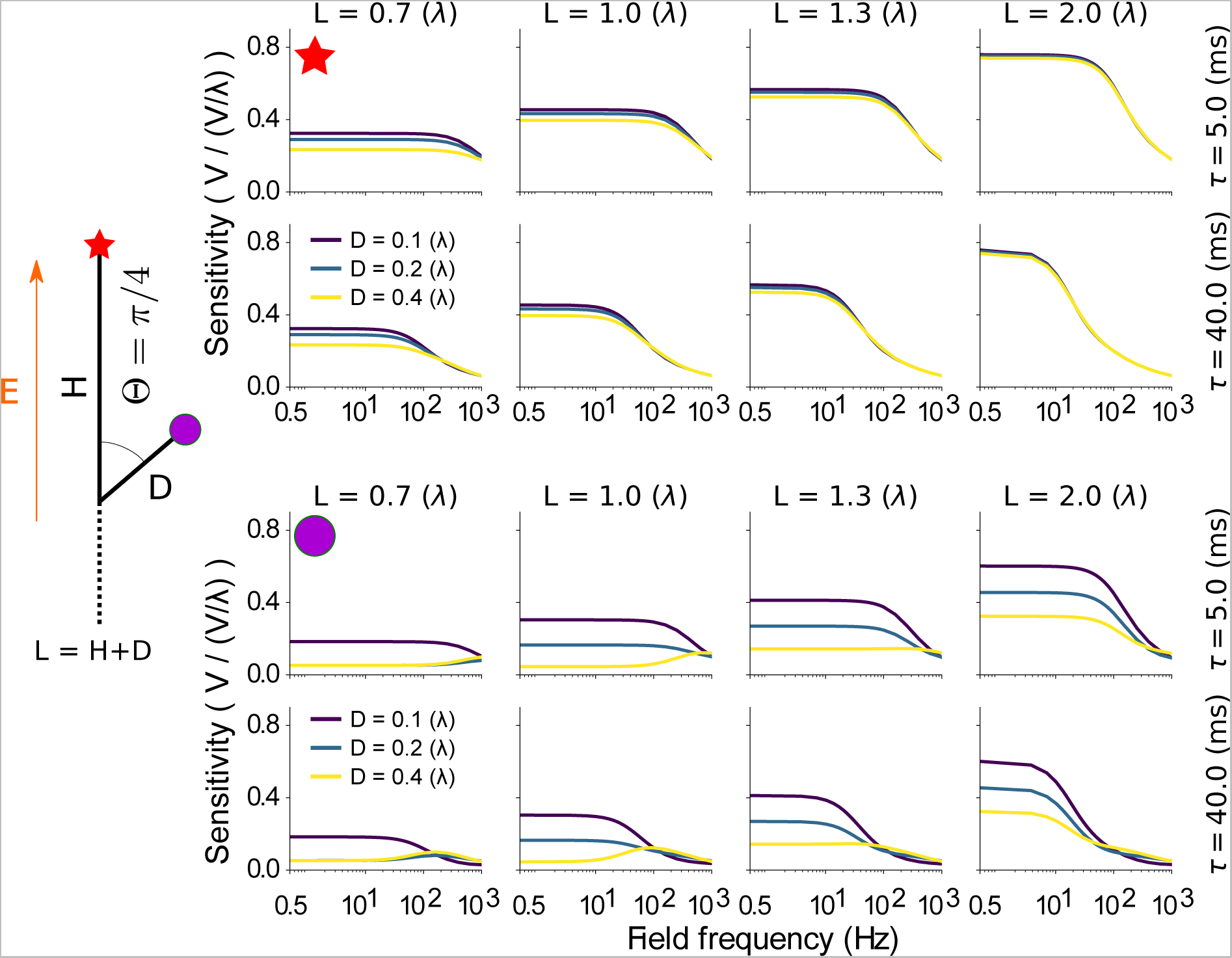
In a passive cable with an acute bending angle, the membrane time constant *τ* determines the resonance frequency. The subplots represent the field sensitivity (in V / (V/λ)) at both cable ends: (top,red star) the unbent branch and (bottom, violet circle) the bent one, as function of the field frequency (x axis). The field sensitivity are displayed for various membrane time constant *τ* (rows for each location), total cable length *L* (increasing from left to right) and bent branch length *D* (color coded). The bending angle is Θ = *π*/4 (rad).

**Figure S5.**
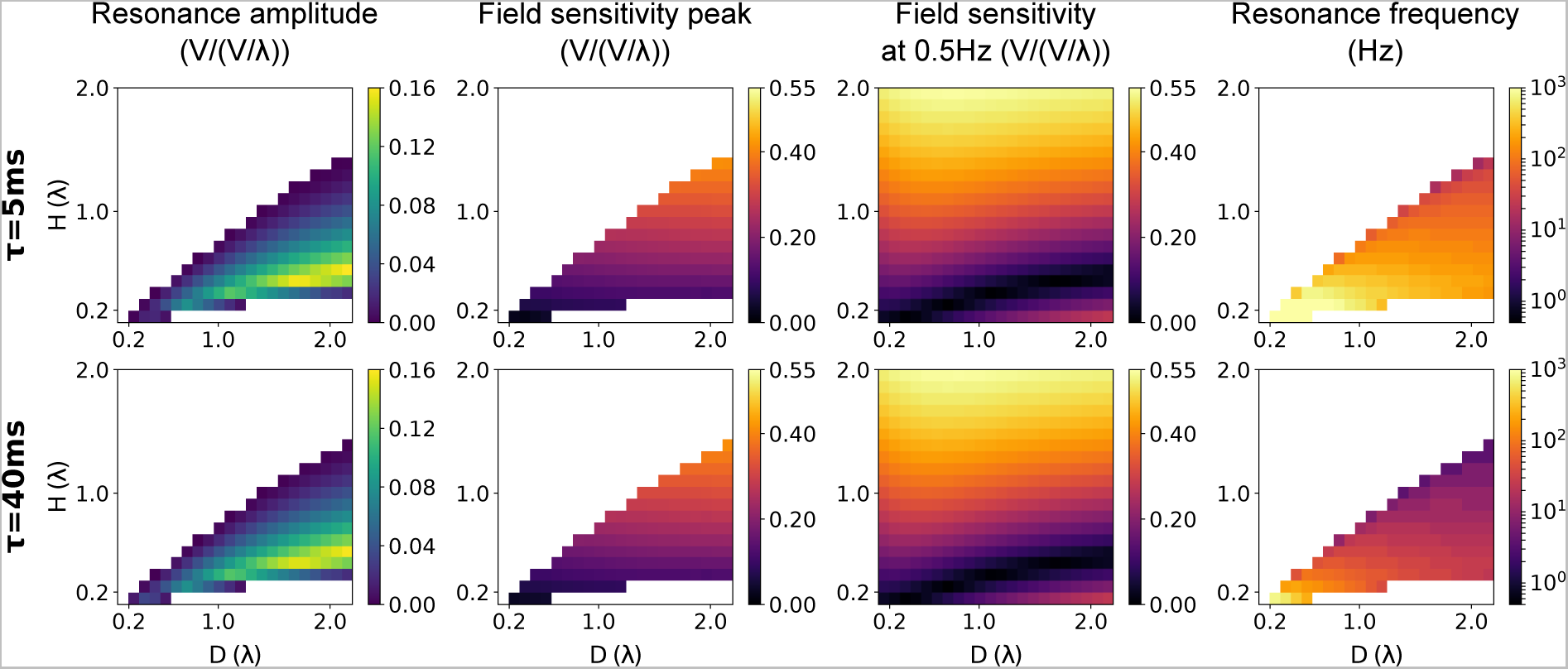
In a passive cable with an acute bending angle, the relative length of the bent branch, *D*, determines the presence of a resonance and the membrane time constant, *τ*, the resonance frequency. We consider the sensitivity to AC fields of a cable with acute bending angle (Θ = *π*/4) measured at the bent extremity. From left to right, the plots represent the resonance amplitude (peak field sensitivity i.e. at the resonance, minus the field sensitivity at 0.5Hz), the peak field sensitivity, the field sensitivity at 0.5 Hz and the resonance frequency. Each of these measures is plotted for various electrotonic lengths of the main branches, H, (vertical axis) and the bent branch electrotonic length, D (horizontal axis). The white areas correspond to the absence of resonance. The rows correspond to different membrane time constants (top: *τ* = 5 ms, bottom: *τ* = 40 ms).

**Figure S6.**
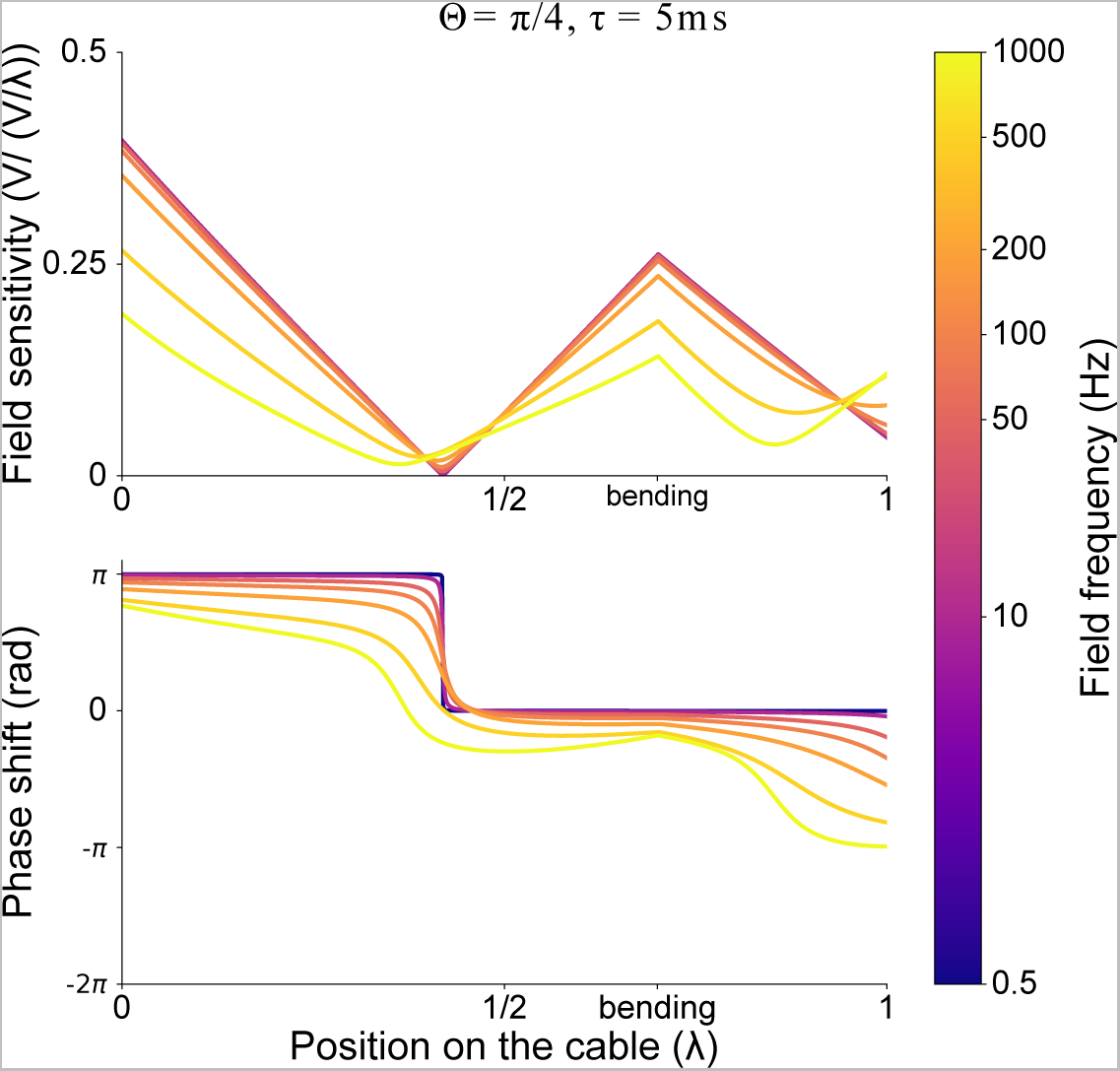
Decreasing the membrane time constant reduces the field sensitivity frequency-dependence of bent cable. Distribution of the sensitivity (top) and phase (bottom) along the bent cable for different field frequencies (0.5, 10, 50, 100, 200, 500 and 1000Hz). Cable parameters are *τ* = 5(*ms*), *L* = 1(λ), Θ = *π*/4 (rad) and *D* = 0.4(λ).

**Figure S7.**
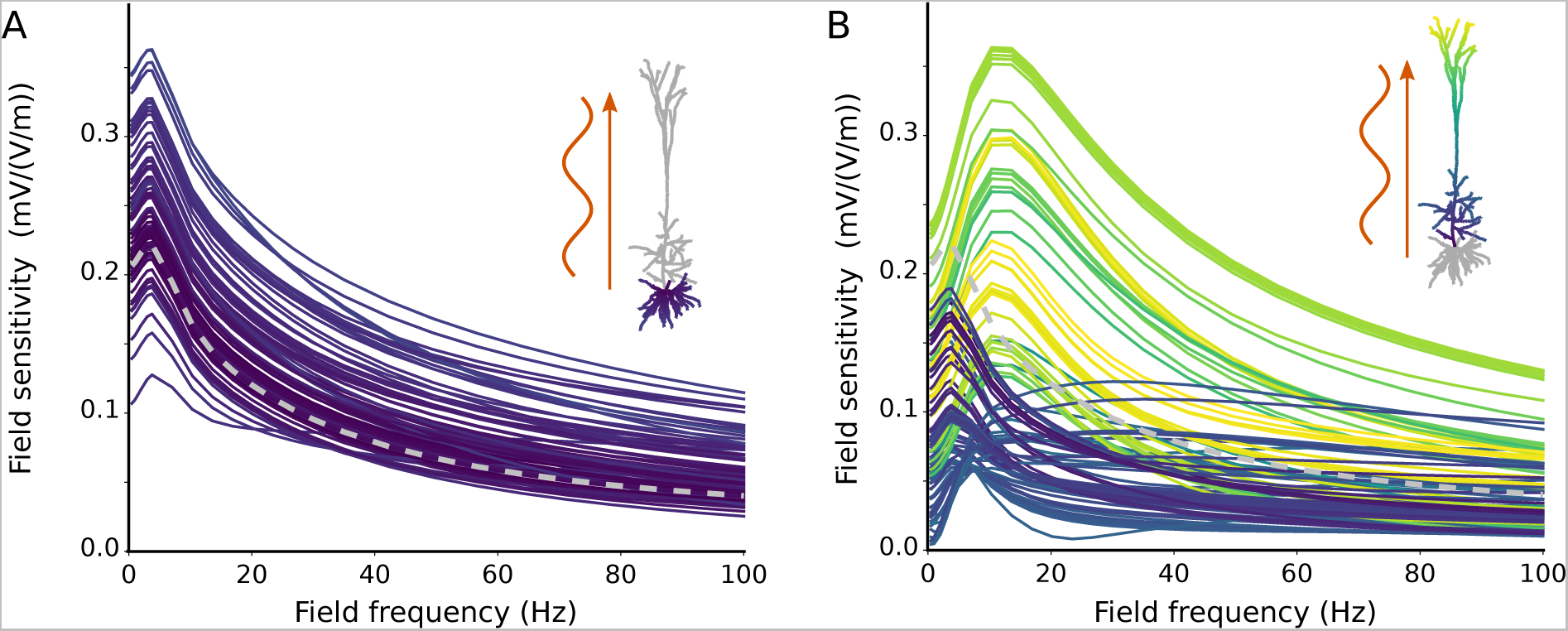
In an active pyramidal cell model, the field sensitivity presents a strong resonance around 10-20Hz at the apical dendrite but not at the soma or the basal dendrite. Frequency-dependent sensitivity of the cell to AC field measured at different locations at the basal (A) or apical (B) dendrites. Colors code the distance to the soma of the considered segment.

**Figure S8.**
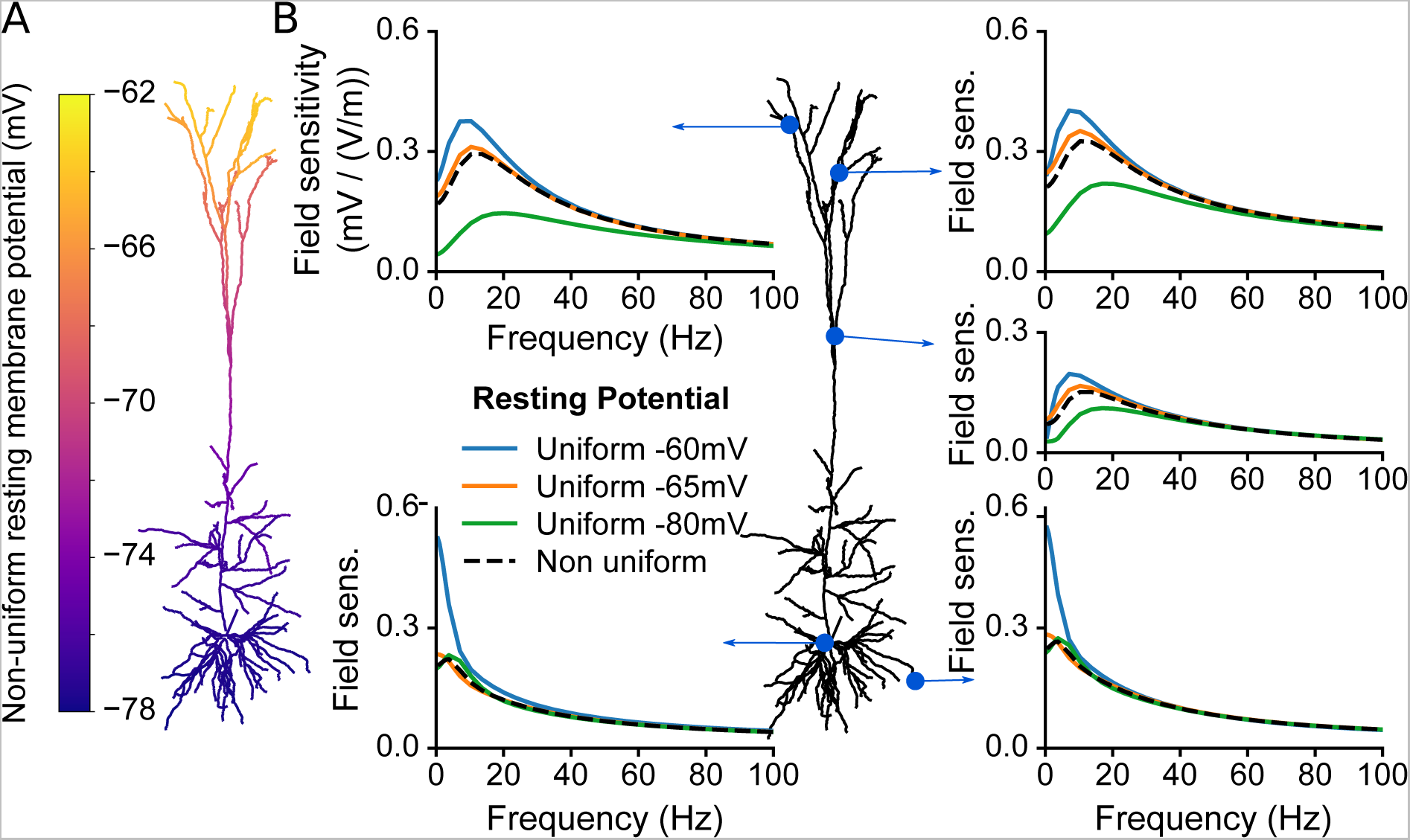
The distribution of the resting membrane potential does not qualitatively affects the field sensitivity profile of the active cell at the apical dendrites. (A) Distribution of the membrane potential at rest, i.e. in absence of electrical fields, in the fully active Hay et al. model. (B) Field sensitivity of the fully active model in case of non-uniform and uniform resting membrane potential. The uniform distribution of the resting membrane was set to an arbitrary value by adjusting the passive leak reversal potential at each dendritic segment independently (see Methods).

**Figure S9.**
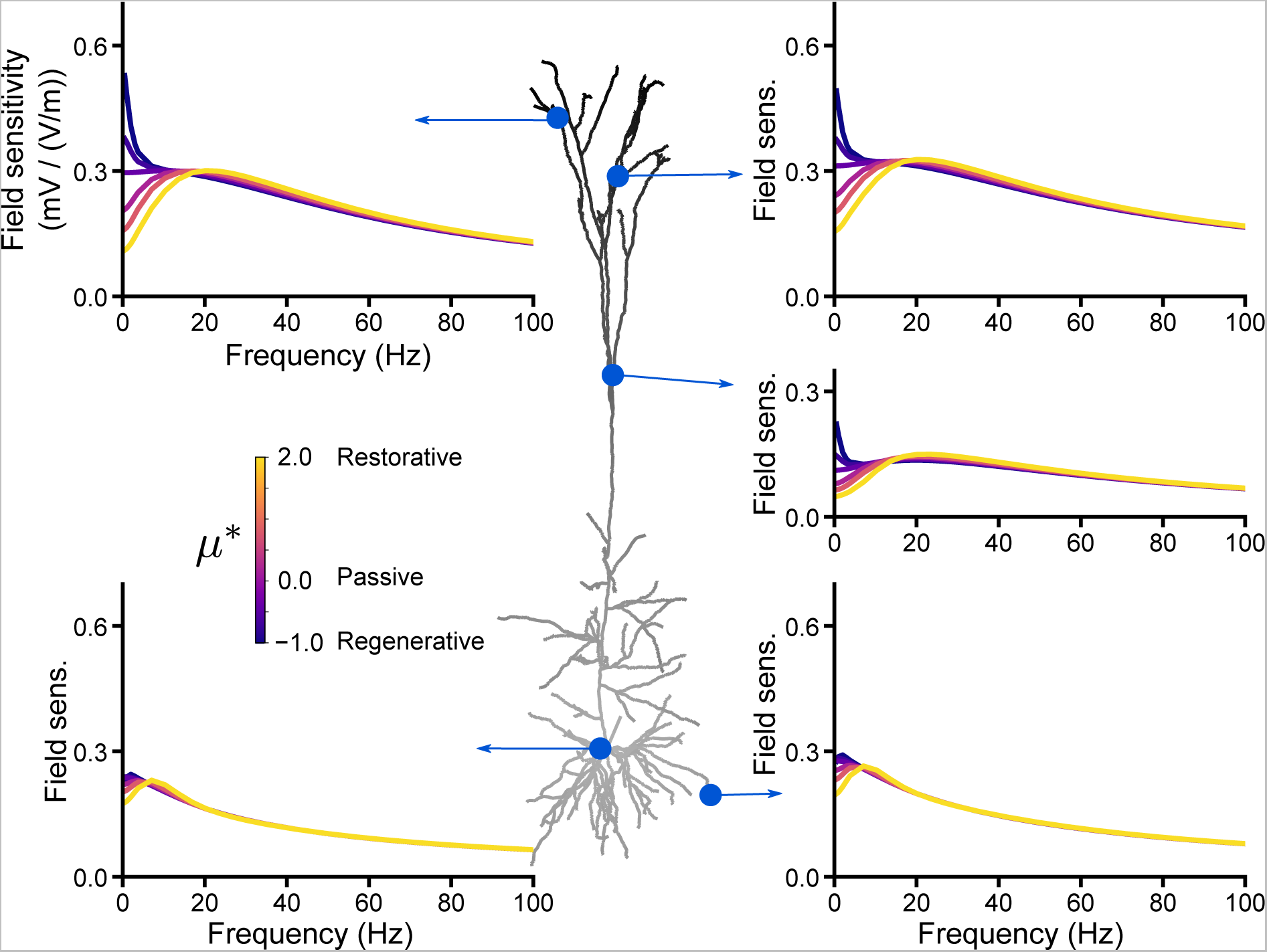
The type of QA channel, with conductance distributed increasingly from the soma, affects more strongly the field sensitivity at apical dendrites than at the soma and basal dendrites. The neuron model includes a leak current and a single QA channel, whose conductance distribution increases linearly with distance from the soma. The shades of grey in the cell plot represent this distribution. *μ*\* determines the type of the QA channel. The plots displays the cell’s sensitivity to AC fields at different locations depending on the values of *μ*\* (color coded).

**Figure S10.**
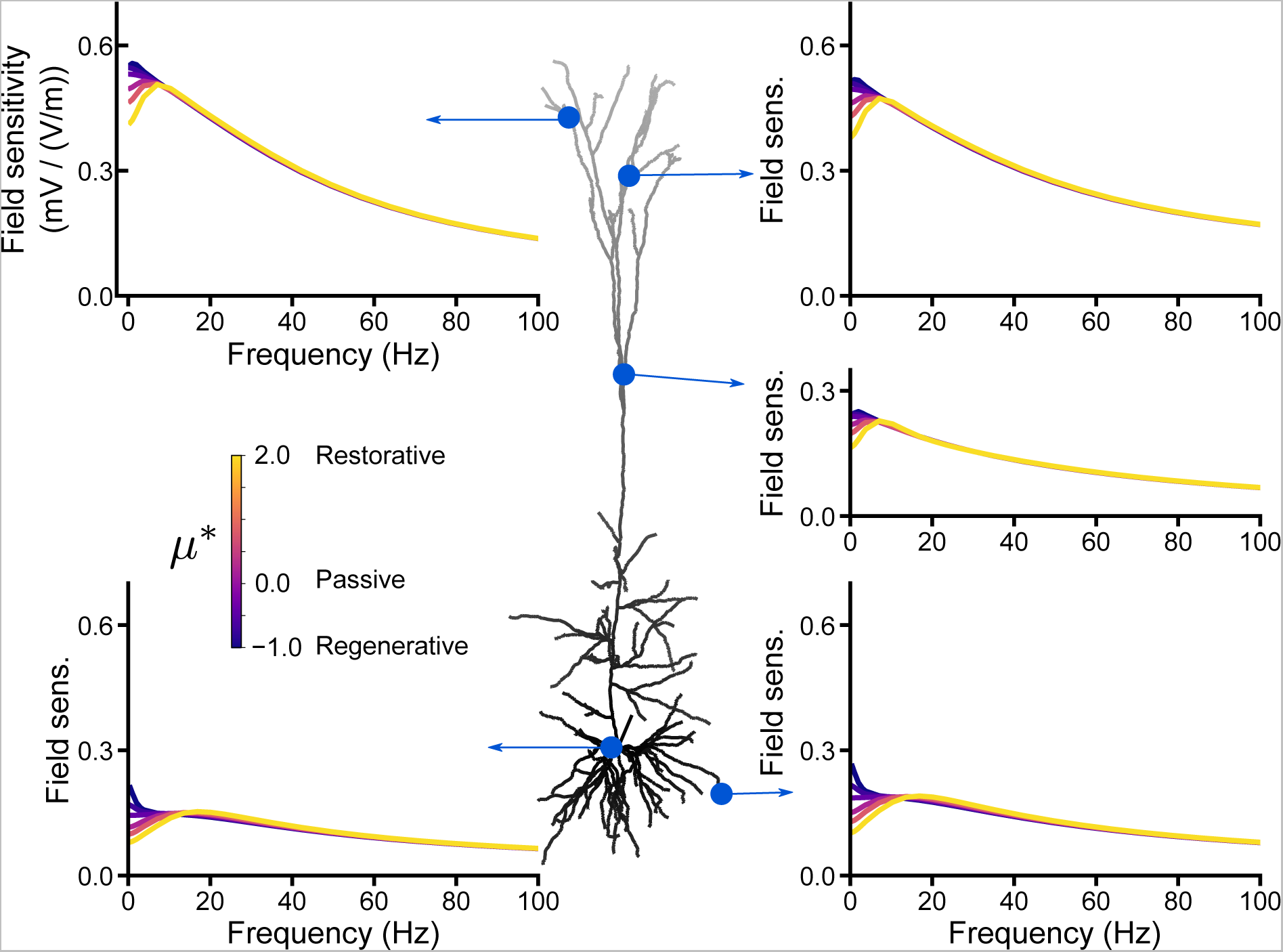
The type of QA channel, with conductance distributed decreasingly from the soma, affects the field sensitivity at the soma, apical and basal dendrites. The neuron model includes a leak current and a single QA channel, whose conductance distribution decreases linearly with distance from the soma. The shades of grey in the cell plot represent this distribution. *μ*\* determines the type of the QA channel. The plots displays the cell’s sensitivity to AC fields at different locations depending on the values of *μ*\* (color coded).

**Figure S11.**
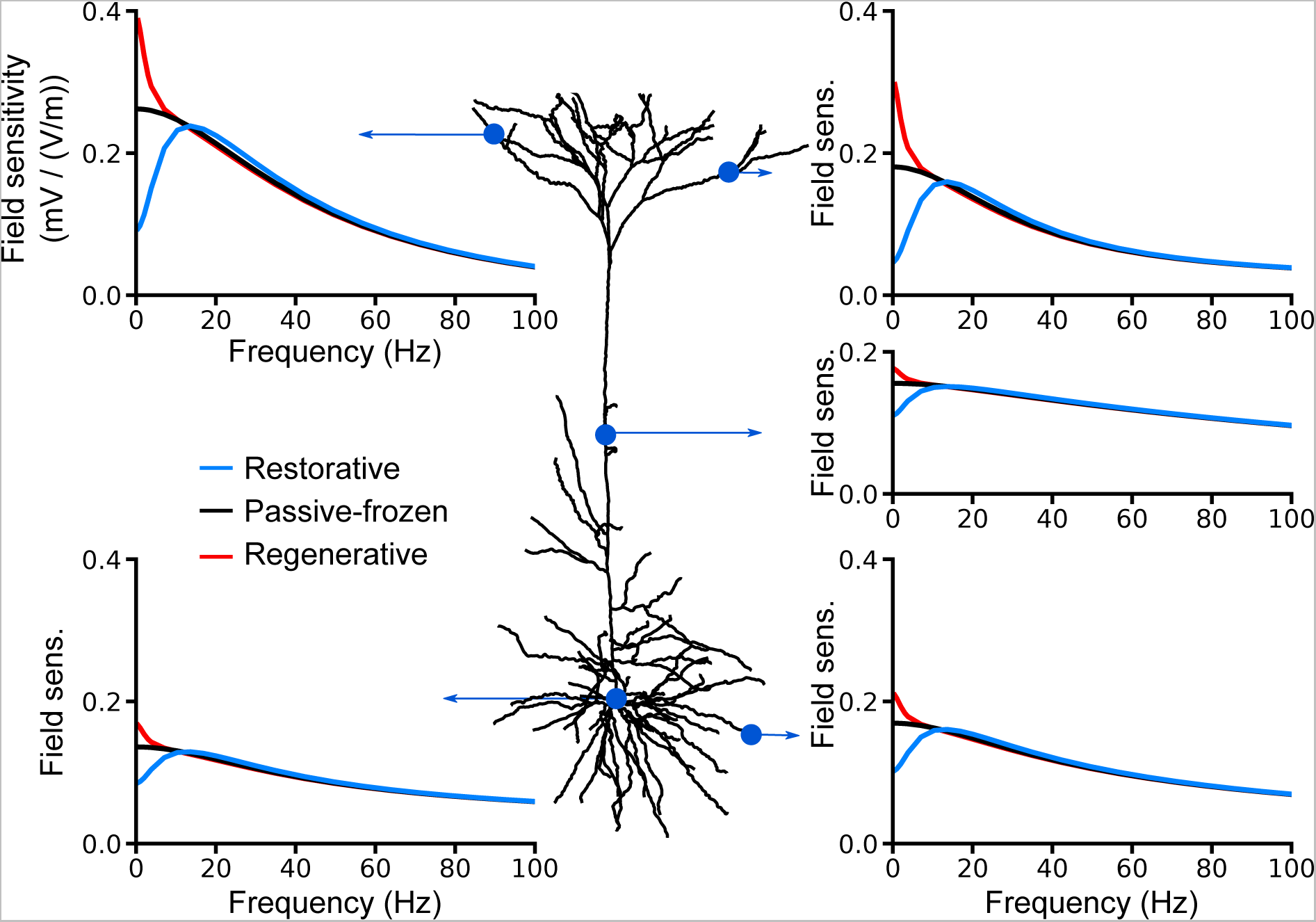
The effects of the channel type on the field sensitivity are transposable to other pyramidal cell morphologies. We consider a neuron model which includes solely a leak conductance and a single uniformly distributed quasi-active channel (QA). We use the reconstructed morphology corresponding to cell 2 in the Hay et al. [16] paper. The plots display the cell’s sensitivity (in mV/(V/m)) to AC fields at different locations in case of restorative (blue, *μ*\* = 2), passive (black, *μ*\* = 0) and regenerative (red, *μ* = −0.5) quasi-active currents.

**Figure S12.**
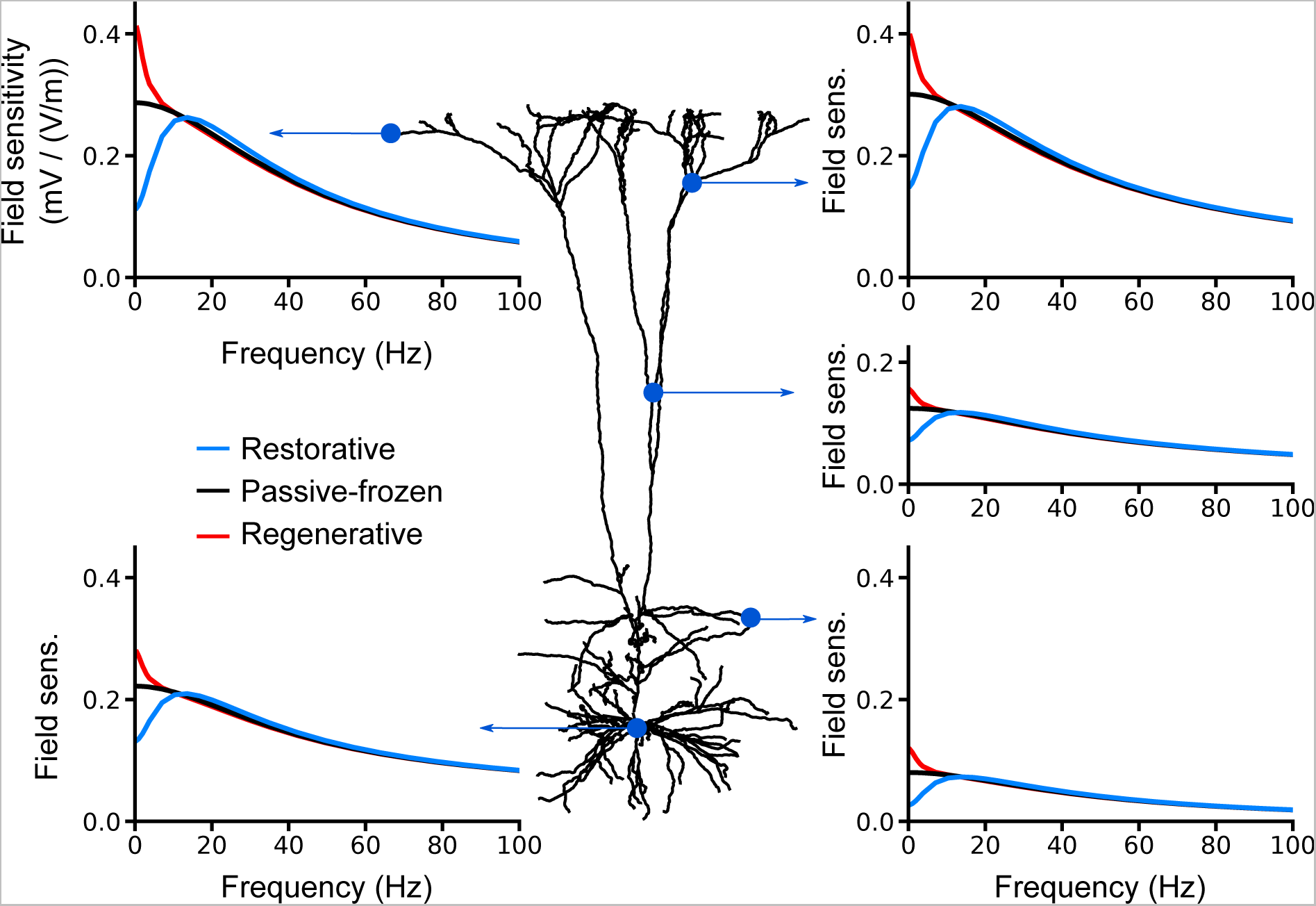
The effects of the channel type on the field sensitivity are transposable to other pyramidal cell morphologies. We consider a neuron model which includes solely a leak conductance and a single uniformly distributed quasi-active channel (QA). We use the reconstructed morphology corresponding to cell 3 in the Hay et al. [16] paper. The plots display the cell’s sensitivity (in mV/(V/m)) to AC fields at different locations in case of restorative (blue, *μ*\* = 2), passive (black, *μ*\* = 0) and regenerative (red, *μ*\* = −0.5) quasi-active currents.

## References

1. Frohlich F, McCormick DA. Endogenous electric fields may guide neocortical network activity. Neuron. 2010;67(1):129–143. doi:10.1016/j.neuron.2010.06.005.

2. Anastassiou CA, Perin R, Markram H, Koch C. Ephaptic coupling of cortical neurons. Nat Neurosci. 2011;14(2):217–223. doi:10.1038/nn.2727.

3. Anastassiou Ca, Perin R, Buzsaki G, Markram H, Koch C. Cell-type- and activity-dependent extracellular correlates of intracellular spiking. J Neurophysiol. 2015;114:608–623. doi:10.1152/jn.00628.2014.

4. Berenyi a, Belluscio M, Mao D, Buzsaki G. Closed-loop control of epilepsy by transcranial electrical stimulation. Science. 2012;337(6095):735–737. doi:10.1126/science.1223154.

5. Marshall L, Helgadóttir H, Mölle M, Born J. Boosting slow oscillations during sleep potentiates memory. Nature. 2006;444(7119):610–3. doi:10.1038/nature05278.

6. Datta A, Bansal V, Diaz J, Patel J, Reato D, Bikson M. Gyri-precise head model of transcranial direct current stimulation: improved spatial focality using a ring electrode versus conventional rectangular pad. Brain Stimul. 2009;2(4):201–207. doi:10.1016/j.brs.2009.03.005.Gyri.

7. Bikson M, Reato D, Rahman A. Cellular and network effects of transcranial direct current stimulation. In: Miniussi C, Paulus W, Rossini PM, editors. Transcranial Brain Stimulation. CRC Press; 2012. p. 55–92.

8. Radman T, Ramos RL, Brumberg JC, Bikson M. Role of cortical cell type and morphology in subthreshold and suprathreshold uniform electric field stimulation in vitro. Brain Stimul. 2009;2(4):215–228. doi:10.1016/j.brs.2009.03.007.

9. Bikson M, Inoue M, Akiyama H, Deans JK, Fox JE, Miyakawa H, et al. Effects of uniform extracellular DC electric fields on excitability in rat hippocampal slices in vitro. J Physiol. 2004;557(Pt 1):175–90. doi:10.1113/jphysiol.2003.055772.

10. Radman T, Su Y, An JH, Parra LC, Bikson M. Spike timing amplifies the effect of electric fields on neurons: implications for endogenous field effects. J Neurosci. 2007;27(11):3030–3036. doi:10.1523/JNEUROSCI.0095-07.2007.

11. Aspart F, Ladenbauer J, Obermayer K. Extending integrate-and-fire model neurons to account for the effects of weak electric fields and input filtering mediated by the dendrite. PLOS Comput Biol. 2016;12(11):1–29. doi:10.1371/journal.pcbi.1005206.

12. Kronberg G, Bridi M, Abel T, Bikson M, Parra LC. Direct current stimulation modulates LTP and LTD: activity dependence and dendritic effects. Brain Stimul. 2017;10(1):51–58. doi:10.1016/j.brs.2016.10.001.

13. Rahman A, Lafon B, Parra LC, Bikson M. Direct current stimulation boosts synaptic gain and cooperativity in vitro. J Physiol. 2017;11:3535–3547. doi:10.1113/JP273005.

14. Deans JK, Powell AD, Jefferys JGR. Sensitivity of coherent oscillations in rat hippocampus to AC electric fields. J Physiol. 2007;583(Pt 2):555–565. doi:10.1113/jphysiol.2007.137711.

15. Major G, Larkum ME, Schiller J. Active properties of neocortical pyramidal neuron dendrites. Annu Rev Neurosci. 2013;36:1–24. doi:10.1146/annurev-neuro-062111-150343.

16. Hay E, Hill S, Schürmann F, Markram H, Segev I. Models of neocortical layer 5b pyramidal cells capturing a wide range of dendritic and perisomatic active properties. PLOS Comput Biol. 2011;7(7). doi:10.1371/journal.pcbi.1002107.

17. Roth BJ, Basser PJ. A model of the stimulation of a nerve fiber by electromagnetic induction. IEEE Trans Biomed Eng. 1990;37(6):588–605. doi:10.1109/10.55662.

18. Koch C. Biophysics of computation: information processing in single neurons. Oxford university press; 1998.

19. Ness TV, Remme MWH, Einevoll GT. Active subthreshold dendritic conductances shape the local field potential. J Physiol. 2016;13(1):1–31. doi:10.1113/JP272022.

20. Koch C. Cable theory in neurons with active, linearized membranes. Biol Cybern. 1984;50(1):15–33–33. doi:10.1007/BF00317936.

21. Remme MWH, Rinzel J. Role of active dendritic conductances in subthreshold input integration. J Comput Neurosci. 2011;31(1):13–30. doi:10.1007/s10827-010-0295-7.

22. Remme MW. Quasi-active approximation of nonlinear dendritic cables. Encyclopedia of Computational Neuroscience. 2015; p. 2553–2556. doi:10.1007/978-1-4614-6675-8 34.

23. Anastassiou CA, Montgomery SM, Barahona M, Buzsaki G, Koch C. The effect of spatially inhomogeneous extracellular electric fields on neurons. J Neurosci. 2010;30(5):1925–1936. doi:10.1523/JNEUROSCI.3635-09.2010.

24. Malik NA. Frequency-dependent response of neurons to oscillating electric fields; 2011. Available from: http://wrap.warwick.ac.uk/55104/.

25. Reato D, Rahman A, Bikson M, Parra LC. Effects of weak transcranial alternating current stimulation on brain activity - a review of known mechanisms from animal studies. Front hum neuroscience. 2013;7:687. doi:10.3389/fnhum.2013.00687.

26. Moore JJ, Ravassard PM, Ho D, Acharya L, Kees AL, Vuong C, et al. Dynamics of cortical dendritic membrane potential and spikes in freely behaving rats. Science. 2017;355(6331):eaaj1497. doi:10.1126/science.aaj1497.

27. Larkum M. A cellular mechanism for cortical associations: an organizing principle for the cerebral cortex. Trends Neurosci. 2012; p. 1–11. doi:10.1016/j.tins.2012.11.006.

28. Carnevale NT, Hines ML. The NEURON book. Cambridge University Press; 2006.

29. Kole MHP. Single Ih channels in pyramidal neuron dendrites: properties, distribution, and impact on action potential output. J Neurosci. 2006;26(6):1677–1687. doi:10.1523/JNEUROSCI.3664-05.2006.

30. Tuckwell HC. Introduction to theoretical neurobiology. vol. 1. Cambridge University Press; 1988.

